# Cortical control of a forelimb prosthesis in mice

**DOI:** 10.1101/2025.09.02.673681

**Authors:** Edouard Ferrand, Zineb Hayatou, Daniel E. Shulz, Maria Makarov, Valérie Ego-Stengel, Luc Estebanez

## Abstract

Robotic upper-limb prostheses aim to restore the autonomy of paralyzed patients and amputees. So far, advances in this field have relied on monkey pre-clinical and human clinical research.

Here, we report on the direct brain control by mice of a miniature mouse forelimb prosthesis. We show that mice implanted with a cortical, microelectrode-based brain-machine interface can learn to control the prosthesis via neuronal operant conditioning, and solve a water collection task in a 2-dimensional and up to a 3-dimensional space. As they learned this task, the mice shaped increasingly consistent prosthesis movements that led to rewards, thanks to coordinated patterns of neuronal activity across the several control dimensions.

Beyond the demonstration of unexpected cognitive and motor control abilities in mice, we anticipate that this preclinical model of upper-limb prosthesis control will be a tool to address several of the most pressing issues in prosthetics controlled by brain-machine interfaces.

## Introduction

Direct brain control on upper-limb prostheses promises a path to restore autonomy for patients with severe motor disfunctions. Patients using an upper-limb prosthesis through an invasive brain-machine interface have achieved control of multiple-joints prostheses (Wodlinger et al., 2015). More recently, a prosthesis control has been obtained in a closed-loop configuration with touch feedback (Flesher et al., 2021). The rodent model has been instrumental in the discovery of these invasive brain-machine interfacing methods (Chapin et al., 1999) and in the exploration of the underlying brain circuitry (Arduin et al., 2013; Clancy et al., 2014; Koralek et al., 2012).

Beyond brain-machine interfacing, mice are an attractive model of the human upper limbs and of their control: mice use coordinated forelimb movements to capture and manipulate their food in ways that resemble some of the primate upper limb behavioral repertoire (Barrett et al., 2020). These movements are encoded in the forelimb area of the primary motor cortex (Barrett et al., 2022), a cortical area that is required for performing forelimb targeted movements (Estebanez et al., 2017; J.-Z. Guo et al., 2015). Sensory-side, in the forelimb area of the primary sensory cortex, representations of forelimb somatosensory inputs have been identified for touch (K. Guo et al., 2020), proprioception (Alonso et al., 2023) and thermoception (Vestergaard et al., 2023). In addition, forelimb embodiment through combined touch and vision has been found in mice (Hayatou et al., 2025). More generally, the self-representation of the body can be studied in the mouse model (Keysers & Michon, 2024; Wada et al., 2016). Further, the unique genetic tooling offered by the mouse model enables an unparalleled dissection of the sensory-motor circuit that underlies the forelimb functional properties (Yamawaki et al., 2021; Yang et al., 2023). These features make mice relevant for the study of the control of upper-limb prosthesis by invasive brain-machine interface. Until now, however, this has been prevented by the lack of limb prostheses that match the specific anatomy of the mouse limbs.

We developed a mouse forelimb prosthesis with 4 degrees of freedom, and we demonstrate its control by mice using an invasive brain-machine interface based on extracellular electrodes implanted in the primary motor cortex (Abbasi et al., 2018, 2023). We show that mice managed to solve a water collection task by learning to structure and coordinate their neuronal activity, and controlling the prosthesis in 2– and 3-dimensional spaces.

## Results

### A forelimb prosthesis for mice

We implanted 7 mice with a 32-channel chronic extracellular electrode in the primary motor cortex (Atlas Neuroengineering, Leuven, Belgium). We then trained the mice to exert direct neuronal control over a forelimb prosthesis, in order to solve a water collection task (Figure 1A) inspired by the ability of water-restricted mice to learn to grab water droplets with their forelimb (Galiñanes et al., 2018). To this end, we designed a miniaturized forelimb prosthesis actuated in closed-loop thanks to the real-time video-tracking of an IR beacon attached to the prosthesis end effector (Figure 1B,C, Supplementary Figure 1A-H). This low-cost, 3D printed prosthesis implemented an elbow as well as a shoulder, for a total of 4 degrees of freedom (Figure 1B). Thanks to a custom robotic controller, the prosthesis movement space (Supplementary Figure 1A) could be restricted to arbitrary subsets of the general control space.

**Figure 1:**
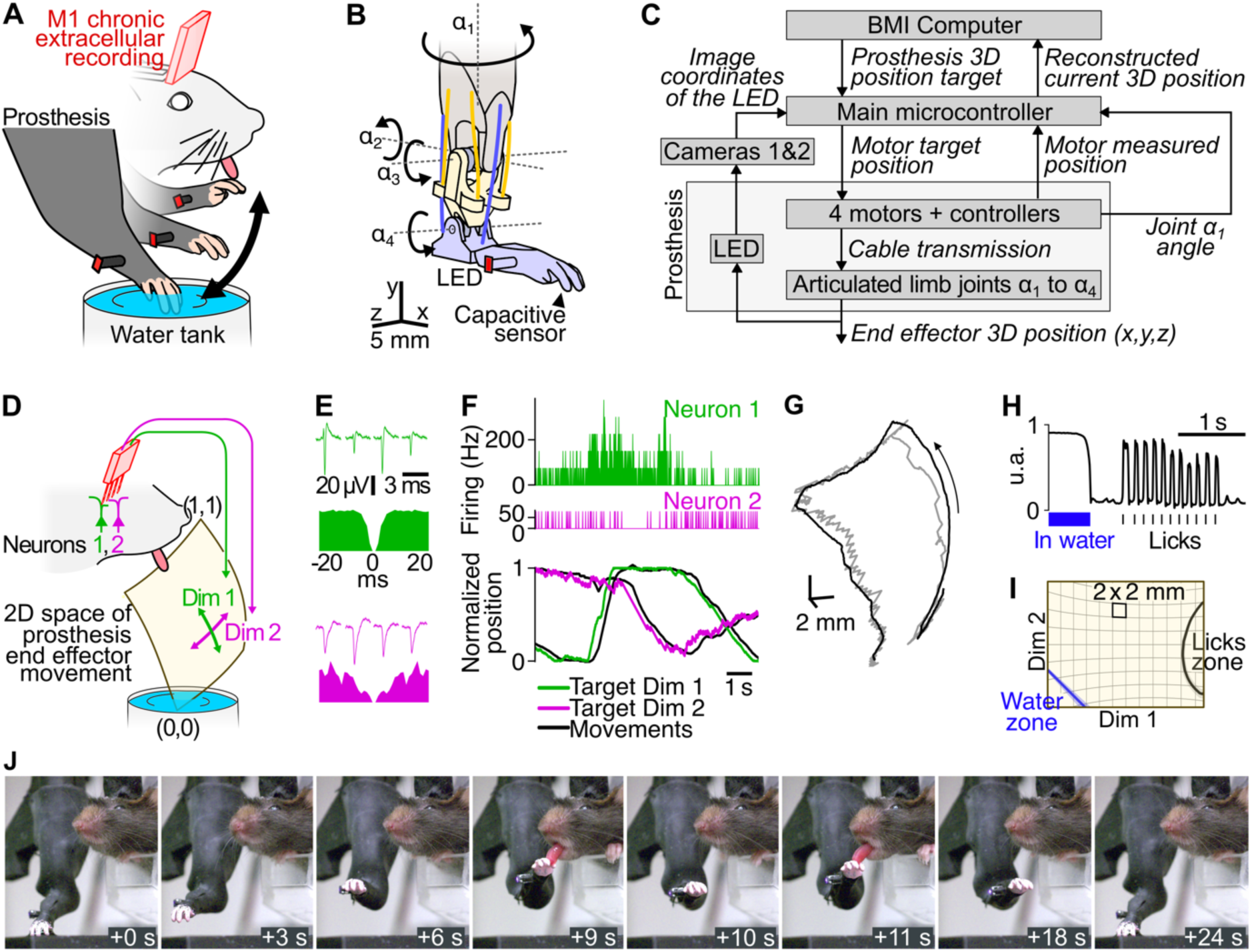
Mice collect water with a forelimb prosthesis controlled by their cortical activity. (**A**) Task design: a head-fixed mouse has continuous control of a forelimb prosthesis via modulation of the activity of individual neurons from the primary motor cortex (M1). To collect water rewards, the mouse must move the prosthesis into the water tank, and then move it close to its mouth to lick the water carried by the prosthesis. (**B**) The 3 moving parts of the skeleton of the prosthesis are articulated through 4 degrees of freedom (DOFs, α_1_ to α_4_). Light colors: skeleton. Saturated lines: corresponding actuation cables. Red dot: infrared LED beacon used to track the prosthesis movements and correct its course. Arrow: location of the capacitive sensor to detect entrance in water and licks. (**C**) Diagram of the prosthesis, control system and its connection to the computer where the brain-machine interfacing is carried out. (**D**) *2*D subspace of control of the prosthesis: one neuron controls displacement over one dimension of the subspace. Coordinates in the subspace range from 0 to 1. (0,0) is under water while (1,0.5) is the closest location to the mouth, where licking for water collection may take place. (**E**) Example of average spiking shapes and autocorrelograms recorded by the tetrodes for the neurons that control the prosthesis displacements. Top: dimension 1 neuron. Bottom: dimension 2 neuron. (**F**) Neuronal activity of the neurons shown in **E**, and corresponding prosthesis movements. Top: for each dimension of control, the spiking activity of a selected, single control neuron is binned by a 13 ms time window and then converted into the normalized coordinate target (bottom) along the corresponding dimension (colored curves). Black: actual position reached by the prosthesis along the dimension. (**G**) 3D view of the target shown in **F**, after converting it into a cartesian target (grey). Black: corresponding prosthesis movements. (**H**) Example of the capacitive sensor readout of individual licks after time spent by the prosthesis in the water tank. (**I**) Population average of the zones in the prosthesis movement space corresponding to the detection of water collection and lick events averaged across all sequences. Background shows a 2 mm grid in the spherical 2D control space, deformed by the projection used for visualization across the article. (**J**) Example of a learned control sequence. The mouse moved the prosthesis from the water tank (0 s), toward its mouth (3 to 6 s after the start), and then licked to collect the water reward (9 s).

In our experiments, we trained the mice to control the prosthesis displacement with their neuronal activity within a 2D space defined in normalized, spherical coordinates. One corner of the space plunged into a water tank, while the middle of an opposite edge was next to mice mouth, where licking actions could collect the water carried by the prosthesis from the water tank (Figure 1D).

The online control of the prosthesis in this 2D space was achieved via a pair of chronically recorded neurons that we selected manually from the multiple single-unit activity recorded by the extracellular electrode (Example in Figure 1E, see Methods). We used their normalized firing rate to control the speed of the prosthesis along one dimension each (Figure 1F). Firing rates above baseline led to positive prosthesis speeds, versus negative speeds for firing below baseline (see Methods, Supplementary Figure 1I). During the task, we observed a good match of the prosthesis displacement to the moving target set by the neuronal activity (Figure 1F,G) while maintaining a spike to movement latencies of 63.5 ms (Supplementary Figure 1F,G) and a movement cutoff frequency of ∼1 Hz (Supplementary Figure 1H).

To ensure accurate detection and timing of (1) the licking of the mice on the prosthesis limb and (2) the entrance of the prosthesis into the water tank, we tracked the readings of a capacitive sensor placed inside the 3D printed “fingers” of the hand end effector of the prosthesis, which is the location where water was captured by capillary forces when plunged into the water tank (Figure 1H). We could then register these events into the prosthesis movement subspace (Figure 1I).

### Mice learn 2D control of the prosthesis to collect water

To initiate the training, we first habituated the mice to the setup for up to 3 days. Their water intake was regulated to build motivation to collect water with the prosthesis, and we initiated daily, 40-minute training sessions on the prosthesis control task. In several mice, we soon observed behavioral sequences (example in Figure 1J) where the prosthesis end effector entered the water tank, then moved towards the mouse mouth. The mouse then licked the prosthesis end effector several times before it came back to the water tank. These action sequences could be detected from the readings of the capacitive sensor located in the prosthesis end effector (Figure 1B,H, Figure 2A), and in particular prosthesis movements from the water tank to a first lick, that we called a Water trip. In addition, because some mice performed multiple licking bursts without going back to the water tank, we defined Rewards as the onset of individual bursts of licks that did not necessarily occur right after a Water trip, but that included licks up to the first 20 after a Water trip, when there was likely still water attached to the prosthesis hand (Figure 2A, Supplementary Figure 1J,K).

**Figure 2:**
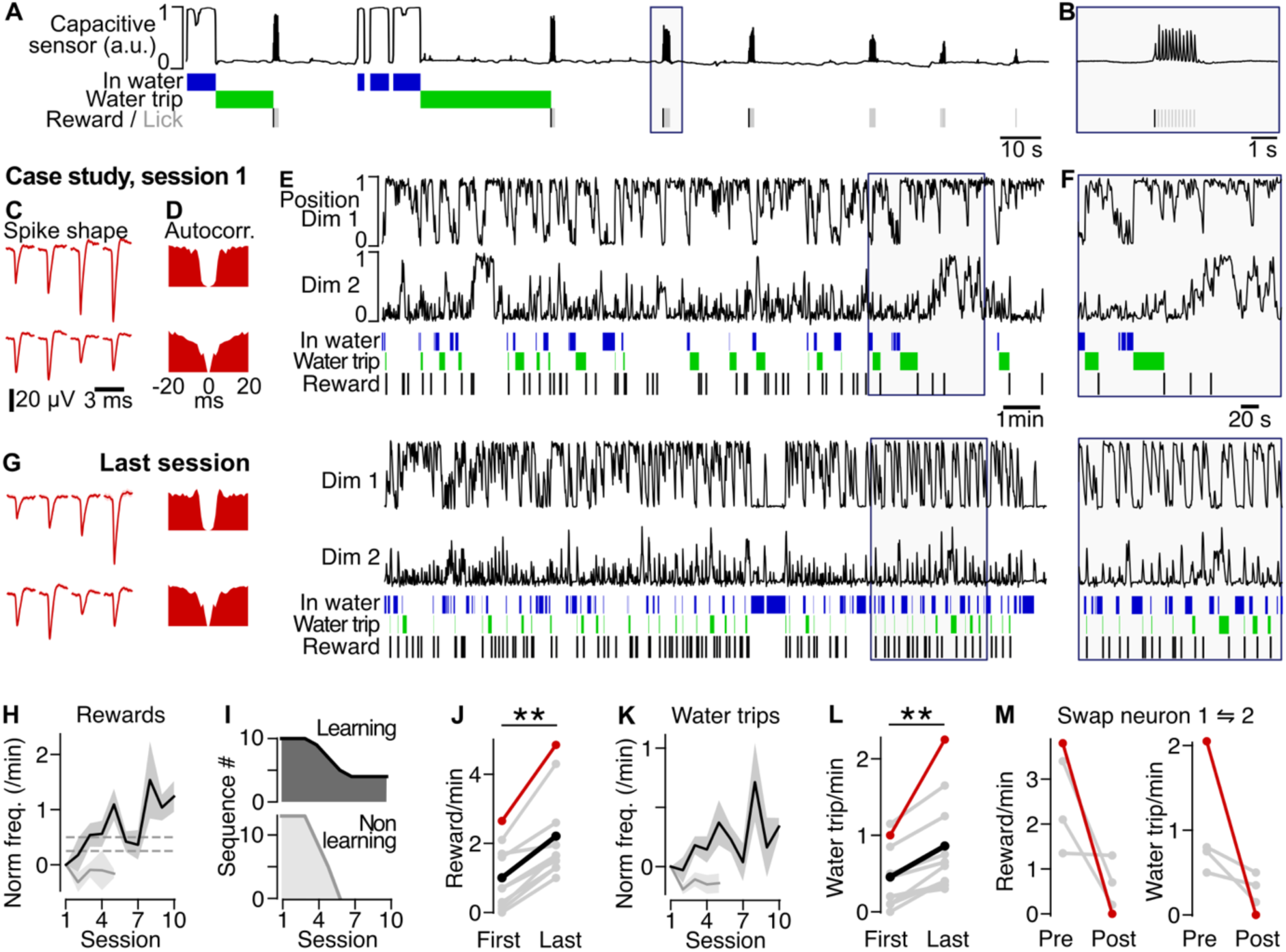
Learning to displace the prosthesis is a 2D space. (**A**) Example reading of the end effector capacitive sensor during the water collection task, corresponding to the time window as in **F**. Top: raw sensor data. Bottom: identification of time spent by the end effector in the water tank (blue, In water). Green: time span of the trip from the water tank to the first following lick. Black: Reward, defined as the first lick of a burst that is less than 20 licks away from the last water exit. Individual licks are shown in gray in the same line. (**B**) Zoom in on **A**. (**C**) In an example mouse, average spiking shapes recorded by the tetrodes for the neurons that control the pros-thesis displacements during the first session of a training sequence. Top 4 spike shapes: dimension 1 neuron. Bottom: dimension 2. (**D**) Autocorrelogram of the neurons shown in **C**. (**E**) Time course of the first session of the example training sequence. Top: movements of the prosthesis in [0-1] space of the two dimensions of the control space. Blue events: times when the prosthesis is in the water tank. Green: travel from the water to a lick (Water trip). Black events: Rewards, defined as lick bursts that include licks among the first 20 after a Water trip. (**F**) Zoom in on **E**. (**G**) Same as **C**-**F** for the last session of the example training sequence. (**H**) Variation of the frequency of rewards during Learning sequences, compared to the first session (dark, 10 se-quences from 6 mice) and Non-learning sequences (grey). Dashed lines are the two threshold levels used to identify Learning sequences. (**I**) Distribution of the duration (number of training sessions) of the Learning and Non-learning sequences. (**J**) Reward frequency in the first versus last session of all Learning sequences (** Wilcoxon test, p = 0.002). Black: average. Red: case study detailed in **C**-**G**. (**K**) Variation of the frequency of Water trips (trips from the water that lead to a lick) during Learning sequences, compared to the first session (dark, 10 sequences from 6 mice) and Non-learning sequences (grey). (**L**) Water trips frequency at first and last session of all Learning sequences (** Wilcoxon test, p = 0.002). (**M**) Left: Reward frequency 20 minutes before (Pre) and after (Post) the neurons in control of dimensions 1 and 2 are swapped. Right: frequency of water trip.

We trained the mice during 17 to 42 days. We focused on sets of consecutive sessions where the recording of the two control neurons was stable (example in Figure 2C-G). We found 23 such sequences (recorded in the 7 trained mice). We analyzed in each session the 20 continuous minutes with the largest reward frequency. In 10 of the identified sequences (from 6 mice), we observed an increase of at least 0.25 rewards/min compared to the first session, followed by a second increase that led to an increase of at least 0.5 rewards/min compared to the first increase (Figure 2H). These sequences with consistent increase in reward frequency across sessions were tagged as “Learning sequences”, while all other sequences were “Non-learning sequences”.

The Learning sequences lasted at least 3, and up to 15 sessions (Figure 2I). Comparing the first and the last session across the population of Learning sequences, we found a significant increase in the reward frequency (Figure 2J), and a significant increase in the water trip frequency (Figure 2K,L).

To test the specificity of the control of the prosthesis achieved by the braibn-machine interface, we permutated in 4 mice the neurons controlling dimension 1 versus dimension 2 at the end of a successful learning session. The high post-training frequency of reward and water trip frequency dropped consistently following the permutation (Figure 2M). This suggests that with training, the mice learned to control the forelimb prosthesis via the brain-machine interface, leading to an increased ability to collect water.

### Stereotyped prosthesis trajectories emerge with learning

As learning progressed, we observed the increasing recurrence of specific prosthesis trajectories, in particular just before lick bursts (including rewarded and non-rewarded lick bursts, example in Figure 3A, see Methods). To analyze and classify these trajectories, we segmented the 2D prosthesis space into a 3×3 grid (Figure 3B) and registered all the 3-step discretized paths followed by the prosthesis across the 9 tiles when leading to a Lick burst (Figure 3C). We found that only a small subset of all possible paths was actually traveled by the prosthesis (on average 11.7 among 60). The most traveled path, that we called Main path, accounted on average for 37.8% of the trajectories that preceded a lick burst, and for more than 30% for 8 of the 10 Learning sequences (Figure 3D-F). The Main paths we identified corresponded to prosthesis trajectories (Figure 3G) that were diverse across mice. Some trajectories crossed a large share of the prosthesis space, enroute from the water tank to the mouth area (Figure 3 Case study 1), while several others were “loopings” close to the mouth, and enabled recurring Lick bursts without repeated visits to the water tank (Figure 3 Case study 2).

**Figure 3:**
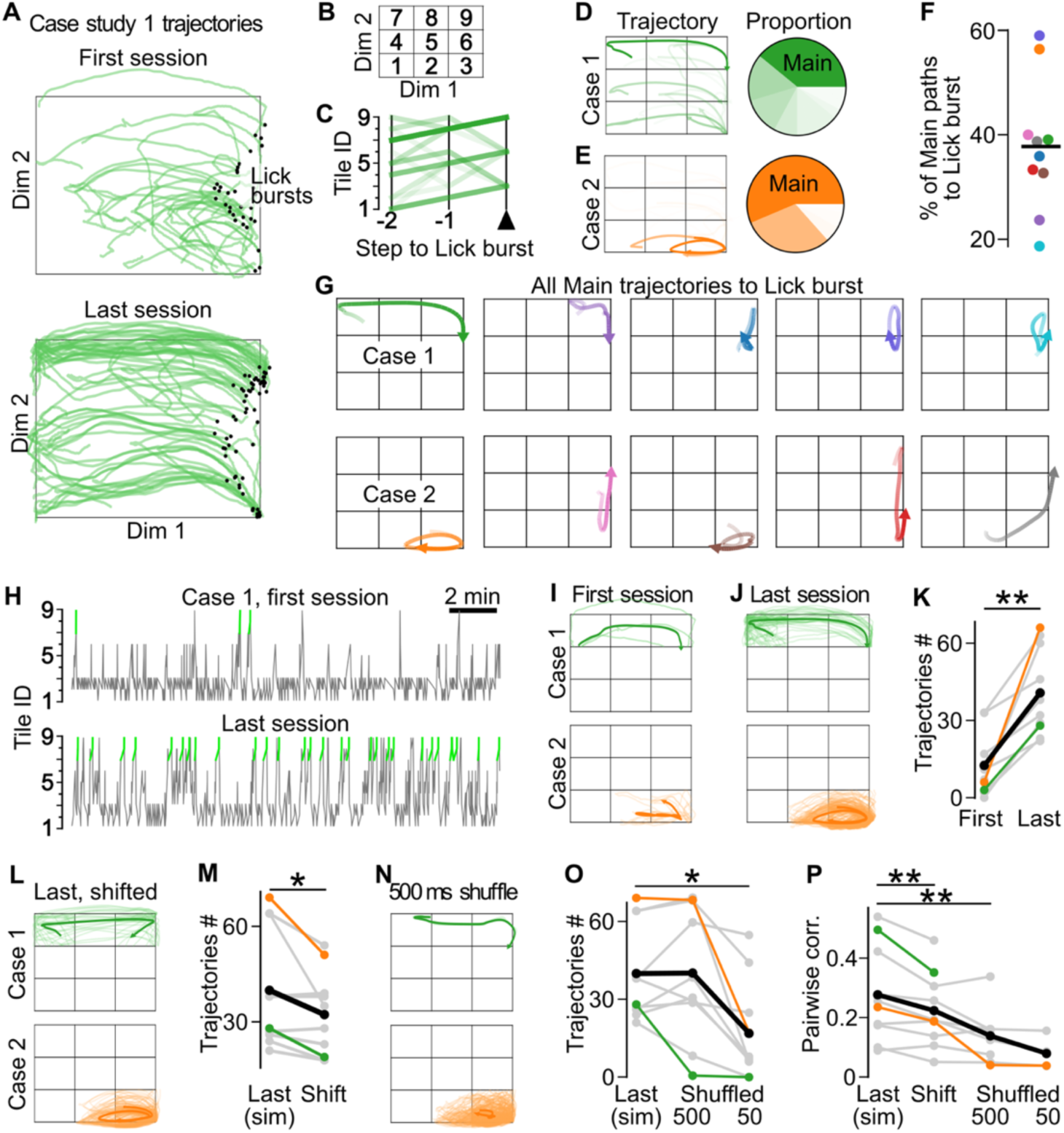
The frequency of the Main trajectory before Lick bursts increases with learning. (**A**) All trajectories 2 s before Lick bursts in the first versus the last session of Case study 1. (**B**) Segmentation of the control space into 9 tiles. (**C**) All series of last 3 tiles visited before Lick burst in **A**. The most frequent path is the “Main path” (7,8,9 in Case study 1). (**D**) Left: average prosthesis trajectories corresponding to the Main and secondary paths (see methods). Transparency is proportional to the proportion of the frequency of a given path. Right: proportion of the different trajectories, shown in the pie chart. (**E**) Same as **D** for Case study 2. (**F**) Proportion of the prosthesis trajectories before Lick burst that follow the Main path, for each Learning sequences. (**G**) Average prosthesis trajectories before Lick burst corresponding to the Main path for each Learning sequences, including Case study 1 and Case study 2. (**H**) Time course of the tiles visited during the first (top) and last (bottom) session of the Learning sequence of Case study 1. Green highlights: Main path. (**I**) Prosthesis trajectories following the Main path (called “Main trajectories”) during the first session, for Case studies 1 & 2. Note that the “Main trajectories” are independent of the Lick bursts. Case study 1 Main trajectories correspond to the green highlights of **H**. Dark: average. (**J**) Same as **I** for the last training session. (**K**) Number of found Main trajectories during the first and last session for all Learning sequences (** Wilcoxon test, p = 0.002). Colored lines correspond to Cases 1 and 2. Black: average. (**L**) Same than **J** but on a simulation of the prosthesis trajectories during the session if the spikes of the neuron controlling Dimension 2 were temporally shifted by +50 s. (**M**) Impact of a shift of Dimension 2 spikes of +50 seconds on the number of Main trajectories (* Wilcoxon test, p = 0.021). (**N**) Same as **L** but with spikes trains shuffled using 500 ms blocks (see methods). (**O**) Number of found Main trajectories after simulations of last session without and with a shuffle with 500 and 50 ms blocks (*: Wilcoxon test with Benjamini-Hochberg correction, p = 0.012). (**P**) Mean pairwise correlation between the prosthesis trajectories predicted from the spikes of the last session, last session shifted, and last session shuffled with a 50 and with a 500 ms block. Correlation was computed only if at least 10 trajectories were found (**: Wilcoxon test p = 0.010 and p = 0.008 respectively).

When tracking the occurrence of the Main path of each Learning sequence from the first and last training session (examples in Figure 3H-J), we found that the number of occurrences of the Main path increased systematically (Figure 3K), suggesting that the trajectories that followed the Main path (called “Main trajectories”) supported the collection of rewards by the mice, and were reinforced by operant conditioning of the task.

To test if these trajectories that we detected in the prosthesis movements could be simply a byproduct of the stochasticity of spike trains, we aimed to disrupt any potential firing patterns that would have been generated by the mice as it attempted to solve the task, and run again our analysis. To this end, we first disrupted any coordination between the two control dimensions of the prosthesis by shifting by +50 s the spike times for the Dimension 2 control neuron versus the Dimension 1 neuron. We computed the resulting prosthesis trajectories by using a neural network-based simulation that converted the spiking activity into prosthesis movements after training on our database of experiments (see Supplementary Figure 2, Methods). We searched in these simulated prostheses movements any occurrence of the Main trajectories (Figure 3L). At the population level, the number of Main trajectories diminished significantly after shifting the spike times (Figure 3M), in comparison to a run of the simulation without the shift (which matched well the actual prosthesis trajectories, Supplementary Figure 2). This suggests that the coordination between the spiking activity of the two control neurons was actively maintained during task performance.

We similarly probed the impact of disrupting the internal time structure of the activity of the control neurons on the occurrence of the 2D Main trajectories. We split the spike trains into blocks that were then randomly permuted to generate shuffled spike trains. This process was carried independently in the two dimensions. We tested shuffling blocks of 500 ms (examples in Figure 3N), and 50 ms, and we used our prosthesis simulation to estimate the resulting trajectories (see Methods). At the population level, the 500 ms shuffle did not impact the number of detected occurrences of the Main trajectories, while a 50 ms block size led to a significantly lower count of Main trajectories (Figure 3O). In addition to the number of detected trajectories, the similarity of the detected trajectory to the average Main trajectory was also reduced significantly, both by the 50 ms shift and even more by the shuffle condition, even with 500 ms blocks (Figure 3P). It is visible when comparing the shape of the trajectories in the control versus shift and versus 500 ms conditions in the 2 case studies (Figure 3J versus 3L versus 3N).

Overall, these observations suggest that the firing modulations underlying the Main trajectories did not stem from spontaneous, random modulation of activity of the control neurons, but rather were deliberate, fast modulations of the neurons’ dynamics that emerged with learning and likely required simultaneous control across both dimensions of the prosthesis movement space.

### Increased coordination of firing rates across control dimensions for main trajectories

To examine the level of coordination between the neurons controlling the 2 dimensions of the task space, we decomposed the 2D Main trajectories leading to a lick burst identified in Figure 3 into their 1D components (Figure 4A). We then searched all the 1D trajectory segments that match the 1D components (Figure 4B,C, see Methods). We carried out the same analysis on the underlying firing activity, leading to the identification of the Main firing patterns underlying the Main trajectory 1D components (Figure 4D) and the identification of matching firing patterns across the whole sessions (Figure 4E,F).

**Figure 4:**
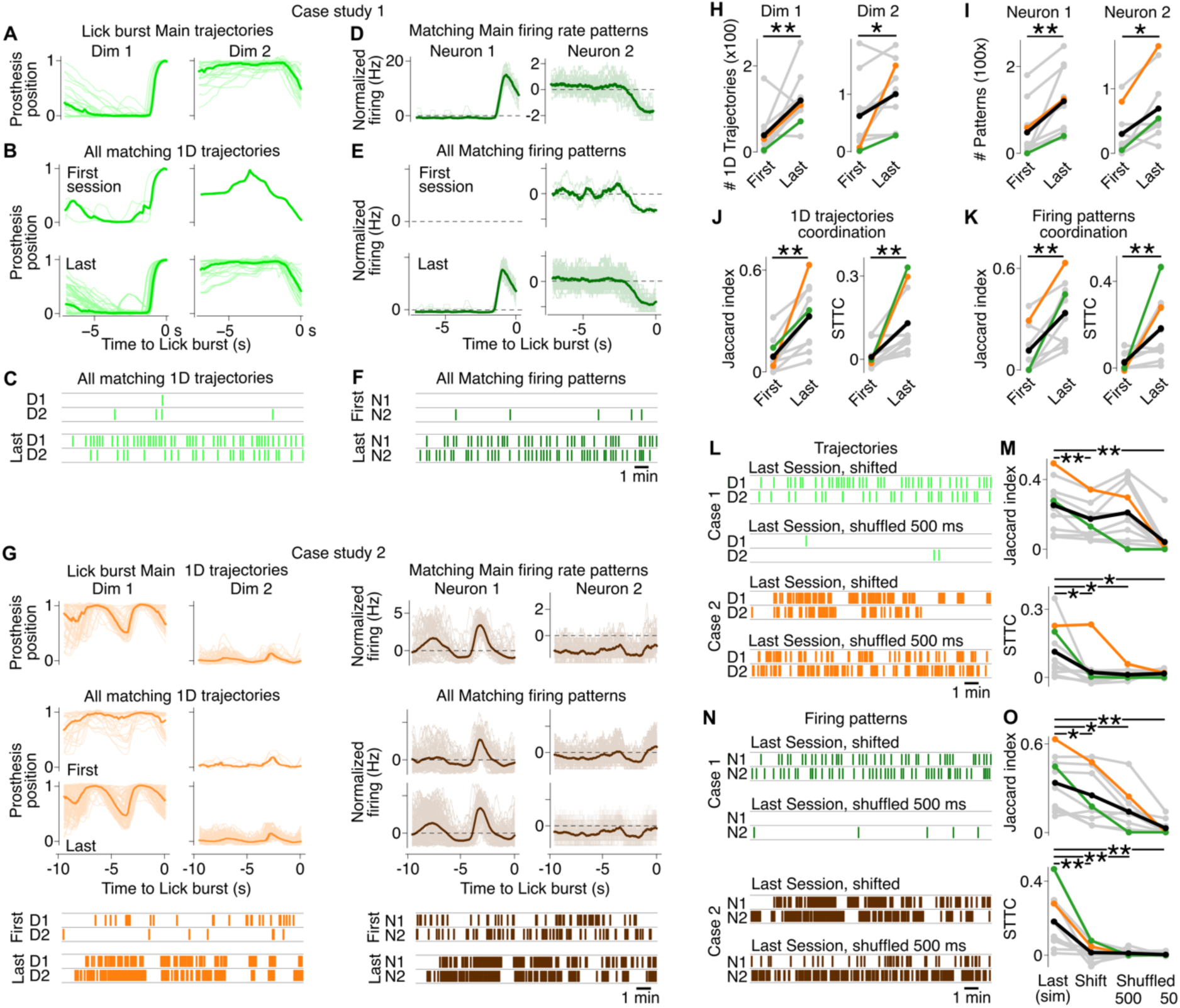
Prosthesis trajectories and underlying activity patterns rely on coordination of the 2 dimensions of control. (**A**) Projection of the Main trajectories leading to a Lick burst on dimensions 1 and 2 of the control space (light green: Case study 1 trajectories). Light color: individual trajectories. Thick dark line: average. All 1D trajectories have been temporally aligned based on correlation to the mean (independently for each dimension). (**B**) “Matching” 1D trajectories found in the first and last sessions of Case study 1. (**C**) Timing of the Matching 1D trajectories displayed in **B**. (**D**-**F**) Same as **A**-**C** for the Main firing rate patterns (dark green: Case study 1 firing) that drive the Main trajectories in Case study 1. (**G**) Same as **A**-**F** for case study 2. Light orange: Case study 2 Main trajectories. Dark brown: Case study 2 firing patterns. (**H**) Number of Matching 1D components of the trajectories detected independently in dimensions 1 and 2 of the control space. Green and Orange: Case studies 1 and 2. Black line: average. Wilcoxon test, *: p = 0.014; **: p = 0.010 (**I**) Same as **H** for firing patterns. Wilcoxon tests, *: p = 0.011; **: p = 0.002 (**J**) Population analysis of the coordination between the Matching 1D trajectories that we detected independently in the two dimensions. Left: Jaccard index. Right: STTC. **: Wilcoxon test, p = 0.002 for both. (**K**) Same as **J** for firing patterns. **: Wilcoxon test, p = 0.006 and p = 0.010 respectively. (**L**) Impact of a 50 s shift (top) and of a 500 ms shuffling (bottom) of neuronal spiking on the occurrence of the 1D Matching trajectories, in Case 1 (green) and in Case 2 (orange). (**M**) Same as **I** for the last session trajectories with no alteration; after a 50 s shift between the two control neurons, and after a 500 or 50 ms shuffling of neuronal spiking. Top: Jaccard index. **: Wilcoxon test with Benjamini-Hochberg correction, p = 0.006 for both. Bottom: STTC. *: Wilcoxon tests with Benjamini-Hochberg correction, from left to right: p = 0.015, p = 0.012 and p = 0.020. (**N**) Same as **L** for the firing patterns underlying the Main trajectories, shifted/shuffled. (**O**) Same as **M** for Main trajectories neuronal activity. Top: Jaccard index. Wilcoxon tests with Benjamini-Hochberg correction, *: p = 0.020 and p = 0.015, **: p = 0.006. Bottom: STTC. **: Wilcoxon tests with Benjamini-Hochberg correction, p = 0.002 for all.

By extending this analysis across all Learning sequences (including case study 2, Figure 4G) we found that the number of 1D trajectory components that matched the components of the Main trajectories increased significantly in both dimensions (Figure 4H). Similarly, both neurons involved in the control of dimension 1 and dimension 2 produced increasing numbers of firing patterns that matched the Main firing patterns (Figure 4I).

We hypothesized that active coordination of the two dimensions of control should result in the synchronized production of the components of the Main trajectories. To test this, we measured the level of coordination between the trajectories/firing patterns identified independently in the two dimensions.

We used a canonical measurement of overlap, the Jaccard index (see Methods), as well as a measurement of event synchrony (applied to the onset of the trajectories/firing patterns) that corrects for the difference in occurrence rate between the two dimensions: the Spike Time Tiling Coefficient (STTC, see Methods). Both the STTC and the Jaccard index revealed a strong and significant increase in the coordination, both of firing patterns and Main trajectory 1D components, from the first to the last session of the learning sequences (Figure 4J,K).

Importantly, both shifting the spike trains in Dimension 1 by 50 s with respect to Dimension 2, and applying 500 or 50 ms block shuffle to the data, diminished significantly the level of coordination measured by the Jaccard and STTC indexes between 1D trajectory components (Figure 4L,M) as well as between the firing rate patterns (Figure 4N,O). These findings further reinforce the view that to solve the prosthesis water collection task, the mice performed coordinated control of a specific prosthesis trajectory in a 2D space.

### Extending the control space to a third dimension

Finally, we asked if the mice may also be able to generate controlled trajectories in a 3D space. To probe this, in a subset of mice, after training to control the prosthesis in the 2D subspace, we extended the control space of the prosthesis to a third dimension. This additional spatial dimension was controlled by a third neuron which used the same conversion to prosthesis movements as the first two dimensions (Figure 5A).

**Figure 5:**
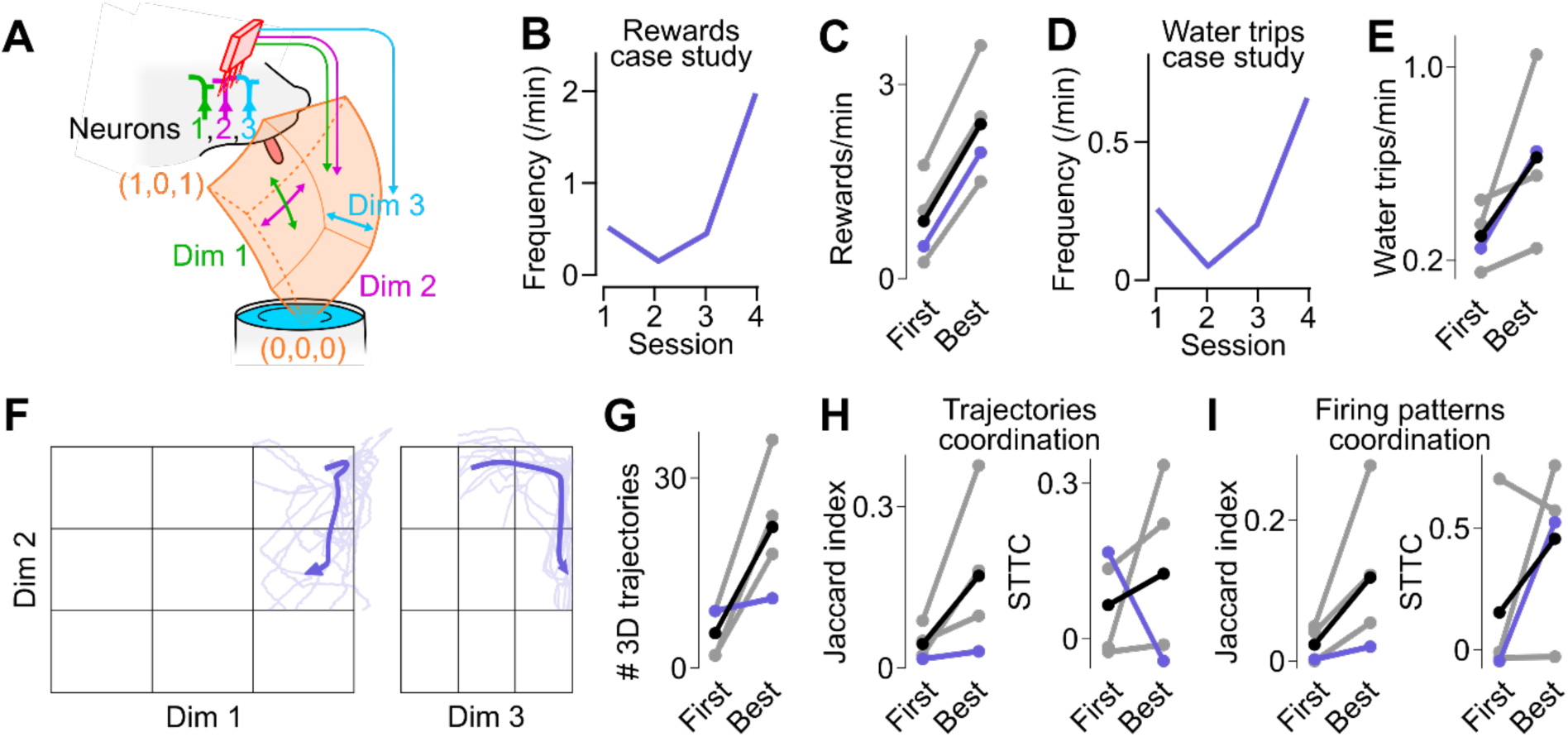
Control of the prosthesis in a 3D space. (**A**) 3D subspace of control of the prosthesis: (0,0,0) is under water while (1,0.5,0.5) is the closest location to collect water. (**B**) Frequency of Rewards across the 3D Learning Sequence in one case study mouse. (**C**) Increase in Reward frequency for all 4 Learning sequences between first and best session. (**D**) Same as B for the Water trip frequency. (**E**) Same as C for the Water trip frequency. (**F**) Average trajectories of the 3D Main paths, shown in two 2D projections of the 3D space. (**G**) Number of 3D trajectories following the Main path in first and best session. Black: average. Purple: case study. (**H**) Change in trajectories coordination between the first and best sessions, measured using a 3D version of the Jaccard index (left) and STTC (right). (**I**) Same as **H** for the Jaccard index.

In this small set of 3D training sessions, to maximize our analysis of control features during high performance sessions, we focused on the best session of the training sequence. In 4 of the training sequences, the mice increased the number of rewards acquired in the 3D space by at least 0.5 rewards/min compared to the first session, and on average by 1.5 rewards/min (Figure 5B,C). This came with a consistent increase in the frequency of water trips (Figure 5D,E). We therefore expected that similarly to the 2D space, the mice would generate stereotyped trajectories just before a lick burst. And indeed, we found that the dominant trajectory (shown for all 3D Learning sequences in Figure 5F) accounted for 20.8% of all trajectories leading to a lick burst. This is a smaller proportion than during the training in a 2D space, but this proportion is drawn from 1258 possible trajectories in the 3D space, versus only 60 in a 2D space.

To probe if the increase in the number of Main trajectories with learning (Figure 5G) is due to coordinated action of the 3 control dimensions together, we carried out the same coordination analysis as previously (see Figure 4) on the generation of Main 3D trajectories from 1D components, asking if they became increasingly synchronous across dimensions. After extracting independently from each dimension the 1D trajectory segments that matched the corresponding components of the Main 3D trajectory, we estimated the amount of coordination of these individual components across the 3 dimensions. From the first to the best performance session, we found a consistent increase in the coordination of the 1D matching trajectories and firing patterns, when measured by a 3D Jaccard index (Figure 5H,I, left). Although more variable, the results of an extension to 3D of the STTC (see Methods) also showed on average an increased synchrony with training (Figure 5H,I, left). Overall, these results suggest that active control of the prosthesis movement patterns can be achieved by mice in a 3D space.

## Discussion

We have shown that mice are able to exert direct neuronal control on a miniature forelimb prosthesis, moving the artificial paw in a 2-dimensional space to obtain water rewards. This space was extended to a third dimension in a more limited set of mice. These findings were possible thanks to the design and fabrication of a forelimb prosthesis for the mouse model, and its connection to the mouse primary motor cortex using a brain-machine interface based on the operant conditioning of individual neurons.

### Physical prostheses are required for pre-clinical applications of brain-machine interfaces

Major advances in motor brain-machine interfacing have relied on feedback on a screen or via sounds (Carmena et al., 2003; Fetz, 1969; Ifft et al., 2013). This is in particular the case in the mouse and rat models (Abbasi et al., 2023; Clancy et al., 2014; Clancy & Mrsic-Flogel, 2021; Goueytes et al., 2019; Prsa et al., 2017). However, when extending the brain-machine interface to a higher number of degrees of freedom, control of such “virtual prostheses” may require a challenging level of cognitive abstraction. In contrast, control of a physical limb prosthesis takes advantage of the familiarity of a subject with its own body movements, as reported by both the visual and tactile sensory modalities. In particular, this multi-sensory input helps support the accurate control of the prosthesis through rich corrective inputs (Flesher et al., 2021; Valle et al., 2025) and may lead to prosthesis embodiment (Risso et al., 2022; Rognini et al., 2019). These benefits of embodied prostheses have led to their use in many brain-machine interfacing experiments in monkeys (Velliste et al., 2008) and in humans (Hochberg et al., 2012). In contrast, in rodent research, physical prostheses have been limited so far to a 1-dimension translation of a reward spout (Arduin et al., 2013, 2014; Chapin et al., 1999). We hypothesize that during our experiments, the ability of the mice to learn to generate control patterns in a 2-dimension and up to 3-dimension space was dependent on the physicality of our prosthesis, and on the rich, physiological sensory inputs made available to the subject, including vision but also whisker contacts on the prosthesis during most phases of its control.

### Robotics for mouse neuroscience

Robotic devices are now a cornerstone of the functional rehabilitation and restoration of limb and body movement and posture, including through the development of advanced robotic upper-limb prostheses (Johannes et al., 2020; Raspopovic et al., 2014; Resnik et al., 2014), and lower-limb and whole-body exoskeletons (Aach et al., 2023; Bach Baunsgaard et al., 2018; Kerdraon et al., 2021). In addition, robotic control methods are increasingly used to directly actuate the body of patients with motor deficits, including the motor control of the lower limbs through stimulations applied to the spinal cord (Wagner et al., 2018) and via direct functional electrical stimulation of the muscles in the legs or upper limbs (del-Ama et al., 2014; Moritz et al., 2008). Beyond rehabilitation, robots and robotic concepts have contributed to better understand the central nervous system functional organization (Ijspeert et al., 2007; Prescott et al., 2024). However, despite the vast investment in the mouse model for biomedical research (Fox, 2007), only limited neurorobotics research has been carried in rodent models so far. Studies have described the development and use of robotic platforms to explore lower-limb mobility restoration in rats (Van Den Brand et al., 2012) and upper limb mobility and motor control in rats (Pasquini et al., 2022; Vigaru et al., 2013) and mice (Alonso et al., 2023), but so far there had been no development of a neuro-prosthetic device for the mouse model.

Here we have designed and implemented a mouse forelimb prosthesis based on Bowden cables mounted on a 3D-printed support skeleton. This approach results in miniaturized actuators that are capable of exerting sizeable force, including surgical robots (Le et al., 2016) and hand prosthesis fingers (Fajardo et al., 2020; Raspopovic et al., 2014). Beyond the 4-joints forelimb structure that we selected, this design strategy is versatile, and may be extended to explore different mouse prostheses, in particular by adding joints, sensors, or by adapting it to obtain a hindlimb prosthesis. We will provide a resource upon publication, containing all required elements to duplicate the prosthesis used in the current project, including the 3D parts plans, a list of all additional mechanical and electronic parts, assembly instructions, as well as the software we developed.

### Extending plasticity-based brain-machine interfacing to multiple dimensions

The first documented cortical brain-machine interface was based on the operant conditioning of individual neurons (Fetz, 1969). Such plasticity-based brain-machine interfaces are immune to representational drift (Sorrell et al., 2021) and require no training of the neuronal decoding stage. This simple design makes them relevant research tools to understand the cortical population dynamics during brain-machine interfacing (Arduin et al., 2013; Clancy et al., 2014) and to study closed-loop brain-machine interfacing (Abbasi et al., 2023; Prsa et al., 2017). With this approach, the arithmetic combination of two (Clancy et al., 2014; Koralek et al., 2012; Prsa et al., 2017) and up to three neurons can be conditioned in mice (Abbasi et al., 2023; Prsa et al., 2017), including to minimize distance of a robot to a reward zone (Prsa et al., 2017). However, speed control — which is used in human studies for versatile motor control (Flesher et al., 2021; Hochberg et al., 2012) has so far only been demonstrated in mice during unidimensional control (Arduin et al., 2013; Goueytes et al., 2022).

In contrast to position control which mainly requires a set of neurons to learn to reach a specific firing rate, speed control demands temporally dynamic modulations of neuronal activity to continuously drive the actuator and stabilize it around the reward zone. Here we show that the plasticity-based brain-machine interfacing approach can be extended to the control of two, and up to three independent axes, under a speed-control paradigm. Mice can therefore learn to generate and coordinate dynamic neural patterns across multiple dimensions. This suggests that plasticity-based interfacing may be relevant to the control of multi-joint limb prostheses by patients, in particular if it is based on the readout of robust signals such as mesoscale cortical readouts (Clancy & Mrsic-Flogel, 2021; Dogadov et al., 2024).

### Fundamental rules of brain-machine interface learning in rodents

Beyond multidimensional control of a physical device, our results suggest that plastic brain-machine learning in rodents may follow the same fundamental rules as in monkeys and humans. In monkeys, brain-machine interface learning has been shown to depend on reinforcing neural patterns consistent with the intrinsic correlations of cortical activity, especially when recording from neuronal ensembles (Carmena et al., 2005; Oby et al., 2025; Sadtler et al., 2014). These constraints on cortical learning (Golub et al., 2018; Oby et al., 2019; Sadtler et al., 2014) provide a stability that is exploited by adaptative decoders (Karpowicz et al., 2025; Orsborn et al., 2014). Similarly, during our experiments in mice, learning did not involve creating entirely novel activity, but rather strengthening existing neural dynamics. Although we did not directly test manifold constraints as in monkey studies (Golub et al., 2018; Oby et al., 2019; Sadtler et al., 2014), our findings are consistent with the view that cortical population structure constrains the learning of brain-machine interfaces. We hypothesize that these constraints account for the diversity of the strategies adopted by the mice, as the specific controlling neurons that were selected — and their functional role within cortical ensembles — likely set the water collection strategy that could be reinforced by individual mice.

In conclusion, beyond the extension of the mouse model into the prosthesis field, our work provides the first evidence for multidimensional brain-machine interface learning in mice. This positions the mouse model to dissect how cortical circuits reorganize to support prosthesis control, and design the next generation of brain-controlled prosthetics.

## Methods

### A forelimb prosthesis for mice

The structure (Umezawa et al., 2022) and general shape (Pyasik et al., 2020; Tieri et al., 2017) of a prosthesis can have an impact on its embodiment. In designing a mouse forelimb prosthesis, we aimed to achieve sufficient similarity with a physiological mouse limb to facilitate the intuitive control of the limb and its embodiment. We therefore designed the shape of the prosthesis and the position of its joints based on a 3D atlas of adult C57BL/6 mice derived from micro-CT sections (Wang et al., 2015). We showed that a static version of this prosthesis (same overall shape, rest position and general material choice) was embodied by mice during a mouse analog of the rubber hand illusion experiment (Hayatou et al., 2025).

Given the miniature size of the forelimb, we chose to articulate only the two largest and most proximal joints: the shoulder, with three degrees of freedom, and the elbow, with one degree of freedom (Supplementary Figure 1B), as these joints are critical to carry out a number of reference motor tasks such as reaching (Georgopoulos et al., 1982) and haptic exploration. The prosthesis mechanical design was structured around a “skeleton” (Figure 1B) that was 3D printed using a Formlabs Form 3B (black resin). It was articulated by miniature ball joints (1 mm inner, 3 mm outer diameters). In particular, at the shoulder, a cardan joint was created by soldering together a set of 4 ball joints. We wrapped the proximal part of the prosthesis, where the joints were implemented, with an outer skin made by molding pre-vulcanized latex (Esprit Composite, France) on a 3D printed shape (Figure 1I), similarly to the design choice made in human-scale prosthetic arms (Johannes et al., 2020). We then applied latex-based modeling paint to mimic the tones of the fur of a C57BL/6 mouse.

The size of the prosthesis led us to operate the prosthesis joints with external actuators through pairs of reciprocal Bowden cables that transmitted forces via the movements of an inner cable (0.2 mm, 316L stainless steel, Filinox, France) sliding inside Teflon tubing (inner diameter 0.3 mm, outer diameter 1.5 mm). The movements were propagating to the prosthesis joints from bipolar stepper motors (NEMA 17) that were fitted with a closed-loop controlled (uStepper S). Motor controllers received their commands from an Arduino-compatible microcontroller (Teensy 4.1) which in turn received a continuous flow of target positions from the brain-machine interface computer (Figure 1B). Thanks to this design, the prosthesis was able to travel in a spherical-like volume (Supplementary Figure 1A). This space was constrained by both mechanical and control design of the prosthesis, reaching 51 mm in horizontal longest dimension and 31 mm in vertical longest dimension.

### Optical tracking of the prosthesis

To ensure the accuracy of the movements of the end effector (the “hand” of the prosthesis), an infrared light beacon (940 nm LED, SFH 4043, OSRAM, Supplementary Figure 1B) located close to the tip of the prosthesis was tracked using a pair of OpenMV machine vision-enabled cameras (Figure 1B, OpenMV H7 R2). Both cameras sent continuously the image coordinates of the LED light blob to the main microcontroller (Figure 1B) at their maximal frequency (around 77 Hz, 13 ms time step). The microcontroller computed the corresponding 3D cartesian position of the LED. This online cartesian 3D tracking required intrinsic and stereo calibrations before the experiment to estimate intrinsic parameters (distortion coefficients and projection matrix) and extrinsic parameters (geometrical transformation between both cameras).

Indeed, the transformation from camera images to cartesian 3D coordinates needed to be accurate and reproducible, both for tracking purposes and to specify target coordinates during motor control. To achieve an overall 50 µm precision, first, the main controller was fed with the intrinsic and extrinsic parameters of both cameras, allowing to track the LED in the cameras coordinate system. Second, we accurately measured the position of the two cameras with respect to the prosthesis. To this aim, we designed a control landmark made of 3 infrared LEDs with a fixed, known geometry (Supplementary Figure 1B). At the start of each use of the prosthesis, their position was sequentially measured with the two cameras, allowing to compensate for any drift in the relative position of the pair of cameras with respect to the prosthesis.

We validated the accuracy of our optical tracking system with a 3-axis micrometric translation (OptoSigma TSD-5-C) that moved with µm precision the LED beacon at regularly-spaced positions on a 20×20×20 mm grid (Supplementary Figure 1C). We found that the camera-based estimate deviated from the actual position by an average of 15 µm between measures within the same startup calibration, and 50 µm between mean measures with different startup calibrations (Supplementary Figure 1D).

### Design of the closed-loop prosthesis control

During prosthesis control, we had access to four pieces of information concerning the skeleton current posture: the 3D coordinates of the end-effector LED, plus the angle α_1_, which could be directly measured by the encoder of motor 1 (Figure 1B,C), as this encoder was deported at the upper extremity of α_1_ axis. Based on this information, the prosthesis microcontroller applied online a reverse geometrical model of the prosthesis to estimate the other 3 current angular positions (α_2_, α_3_ and α_4_) with a unique solution under the known constraints of the prosthesis control. These constraints were (1) limits on the angle values and (2) the linked behavior of α_1_ and α_2_ to reach the BMI target: during the conversion from a cartesian target to an angular target, α_2_ was set proportionally to the z target and α_1_ was then set to complement for the α_2_ contribution on z in order to reach the z target.

After receiving a cartesian command from the control computer, the main micro-controller converted this command to an angular command, and measured the current distance between the goal and estimated angular positions. This distance was then converted to motor displacement commands through proportional gains to update the position of all 4 motors. This closed-loop process was done following each update from the LED tracking system, thereby continuously minimizing the distance between the target position commands issued by the BMI computer and the actual position of the prosthesis.

### Prosthesis benchmarking

Taking advantage of the accurate measurement of the 3D position of the prosthesis, it could achieve static positioning at fixed targets with a variance that was lower than 50 µm (Supplementary Figure 1E). Regarding dynamical positioning, on average the prosthesis responded to a step displacement command of 10 mm in each cartesian dimension with a latency of 63.5 ms (Supplementary Figure 1F,G). Finally, we probed the ability of the prosthesis to reproduce movements with a rich frequency content. A frequency response analysis from sinusoidal inputs (Bode plots, Supplementary Figure 1H) showed that the prosthesis was able to track accurately movements commands up to approximately 1 Hz (Estimation of cutoff frequencies at –3 dB from the plots: x axis 1.5 Hz, y axis 1.8 Hz and z axis 0.85 Hz).

In addition to the position tracking, we also included in the prosthesis “hand” a capacitive sensor, by soldering a shielded conducting cable to a capacitive sensor (Balluff BCS001A) located on the inside the three biggest fingers. This provided straightforward timing of the contacts of the prosthesis with the water inside the tank, as well as of the licks of the mouse on the prosthesis (Figure 1G).

### Replicability of the prosthesis

We designed our prosthesis with the aim of being an affordable device that could be designed with a standard 3D resin printer (Formlab 3B). Resources that are required to assemble the prosthesis will be detailed online upon publication of the article. In particular, we will provide a parts list as well as the code implemented in the control computer, the microcontroller, the motor controllers and the camera controllers, as well as 3D files for the prosthesis parts, and assembly directions. So far, we have replicated the prosthesis 3 times within the laboratory, and one was achieved in an external laboratory (Alex Caldas, ESME Sudria engineering school, France).

### Mouse experiments

All animal experiments were performed according to European and French law as well as CNRS guidelines, and were approved by the French Ministry for Research (Ethical Committee 59, authorization 25932-2020060813556163v7). We minimize the number of mice generated in our institute, we used surplus mice from lines without specific phenotype, and which matched our experimental needs. Specifically, 3 EMX1-Cre x Ai95 and 4 EMX1-Cre female mice were used.

The mice were implanted between P56 and P65 with a titanium head-fixation post (Ymetry ITHB) and a 32-channel extracellular silicon probe (Atlas Neuro E32T8+R-100-S4-L6 NT). Implantation surgeries were carried under Isoflurane anesthesia (4% for induction and 1-1.5% for maintenance) on a heated pad, while the mouse was held by a nose clamp. After a subcutaneous injection of lidocaine (4 mg/kg) the scalp was removed, the conjunctive tissue resected and the exposed skull was cleaned and coated with cyanoacrylate glue. The head-fixation post was glued with Cyanoacrylate glue, and a first layer of dental cement (Lang Dental, Jet dental cement) was applied, while keeping the area of the implantation of the electrodes free of cement on the left hemisphere. A ground screw was implanted above the right hemisphere, and a craniotomy was performed above the forelimb primary motor cortex area in the left hemisphere (+0.5 mm anterior–posterior and +1.7 mm mediolateral relative to bregma). We then lowered the extracellular electrode down to cortical layer 5 (tip of the electrode at 800 µm deep in the cortex). Following electrode insertion, a low-toxicity silicone sealant (Kwik-cast, WPI) was used to fill the craniotomy. Finally, we coated the entire assembly with additional layers of dental cement. Mice received a subcutaneous injection of anti-inflammatory medication (Meloxicam, 1-8 mg/kg) and were monitored during their recovery in a temperature-regulated cage.

### Spike sorting, control algorithm and normalized firing rates

Spikes were recorded using a Cerebus Neural Signals Processor (Blackrock Neurotech, Salt Lake City, USA), and sorted in an amplitude feature space with the Cerbus software suite, before further online processing with our inhouse C++ software.

We chose units with large waveforms (putative regular-spiking neuron) and sufficiently high firing rates (examples in Figure 2A-E). Each selected unit was attributed to one dimension of control.

At each time step dt (every 13 ms, synchronous with camera tracking), spikes of each unit that fell within the ongoing time bin [t, t+dt[were counted and linearly converted into a speed along the dimension controlled by that unit (Supplementary Figure 1I):

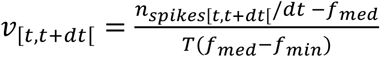

where *f_min_* is the 1^st^ percentile firing rate and *f_med_* is the median firing rate measured during a calibration period of 3 minutes before the ongoing session (after removing all bins with no spike) and T is a time constant controlling the slope of the linear function linking the frequency to the speed (see Supplementary Figure 1I). This parameter can be interpreted as the required amount of time for the target to move from 1 to 0 on the controlled dimension if the frequency is maintained constant at *f_min_* (Dim 1: T = 5 s, Dim 2: T = 4 s, Dim 3: T = 4 s).

The target position *p_t+dt_* along one dimension was calculated from the previous position *p_t_* by using this speed, followed by clipping between 0 and 1:

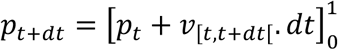

The resulting target position vector within the 2 or 3 Dimensions of control was then converted into cartesian 3D coordinates sent to the prosthesis controller (Figure 1F).

### Task training

After 1 week minimum of recovery, all mice performed 1 session of 40 minutes each day. Before training, mice underwent habituation sessions over 3 days: 1 passive session on the setup while recording spiking activity, followed by 2 sessions during which the prosthesis moved according to predefined movements along Dimension 1 (Supplementary Figure 1J), and the mice learned to lick the water carried by the prosthesis. During habituation and training, a mirror was positioned on one side of the mouse to increase the amount of visual feedback from the prosthesis. After the initial training, 2 sessions were sometimes realized in one day when necessary to perform a particular control experiment. During the entire behavioral protocol, the mice received oral Metacam to limit the inflammatory response to the implanted electrodes, which can deteriorate the quality of recorded signals.

### Capacitive sensor-derived definition of the water and licks zone in control space

The capacitive sensor raw signal was recorded through the Blackrock Cerebus acquisition system (Figure 1G). The signal was analyzed offline to detect both the entrances in the water tank, and tongue contacts. To discriminate those two categories of contact, we used the prosthesis position, setting a positional threshold on Dimension 1: if Dimension 1 position was greater than 0.7, a peak increase of the normalized sensor signal of 0.15 between maximal and both pre and post minimal values within a 200 ms windows was detected as a lick. The licking time was considered as the time of the maximal positive derivative value of the signal. If Dimension 1 position was lower than the threshold and the normalized signal crossed a 0.3 threshold, the corresponding times were considered as a water entry or exit depending of the direction. All contacts were then filtered to only retain contacts separated by a minimal time interval (500 ms for a water entry / exit and 100 ms for licks). Finally, the times measured with the capacitive sensor signal were corrected with an affine transform to account for the distinct clocks of the main computer and the Blackrock recording system.

From these analyses of the capacitive sensor, for each Learning Sequence, we defined a water zone and a licks zone inside the 2D control space of the prosthesis. The borders of the water zone were defined as the locations of the prosthesis at the time when the capacitive sensor signaled the prosthesis entrance in the water tank. A linear regression was used on all the water entries location of a Learning Sequence to define the exact water tank entrance threshold from the X,Y prosthesis position pairs acquired through the capacitive sensor. The licks zone was defined as the smallest 2D convex area containing every location of the prosthesis at the exact time of the licks performed by the mouse during the learning sequence (its convex hull).

For the zone representation of Figure 1.H, we computed the average zone of the water zone as the segment between the averages of the intersection points of the edges of the control space with the water zone edge lines computed from individual Learning Sequences. We also computed an average lick zone as the average of the 2D convex hull of the lick zones identified across Learning Sequences. To this end, we carried a ray tracing approach: we computed the barycenter of all licks across all sessions/sequences, and we projected multiple rays from this central point. We computed their intersection with each the licks convex hull found across Learning Sequences, and derived a ray-by-ray average lick zone.

### Definition of Rewards and Water trips

Using the timing of licks and of the exits from the water tank provided by the capacitive sensor, we identified Water trips and Reward events. We defined Water trips as the time windows from the exit of the prosthesis off the water tank, up to the first following lick, and we defined a Reward as the onset of a lick burst (a lick sequence with intervals lower than 500 ms) that included at least one lick that delivered water to the mice. Indeed, in our experiments, some mice licked repeatedly the prosthesis hand without frequent water trips, to the point where the mouse licked a prosthesis that did not hold any water. We therefore aimed to determine the number of licks after a Water trip that deliver water to the mouse, and only count as a Reward a lick burst that contained water-delivering licks.

To carry this estimation, we used the habituation sessions, where a predefined pattern of prosthesis movements cycled from the water tank to the lick area (Supplementary Figure 1J). Once the prosthesis arrived at the lick area, the mice had ample time (several seconds) to lick freely the collected water before it would automatically go back to the water tank. We used the number of licks performed by the mice during this period as an indicator of the number of licks that are required to drink all the water carried by the prosthesis. We built a distribution of the number of licks per prosthesis cycle in 4 habituation sessions from 2 mice (Supplementary Figure 1K). We chose the 95^th^ percentile of this distribution, corresponding to 20 licks, as a conservative threshold to determine if a lick actually provided water to the mouse. Therefore, all licks following a Water trip until the 20^th^ were considered as providing water to the mouse, and the onset of a lick burst was registered as a Reward if it contained such a lick.

Finally, we aimed to control if this 20-lick threshold was meaningful. To this end, we measured in 2 mice the amount of water left on the prosthesis after pluging the prosthesis in the water tank and allowing the mouse to perform repeated licking on the prosthesis. The licking (we counted individual licks with the capacitive sensor) was interrupted by the rapid retractation of the prosthesis away from the mouse, and its end effector was throughly whiped with a piece of hygroscopic paper towel that was weighted on a precision scale. Water evaporation was stopped by immediate enclosure of the paper samples in Eppendorf tubes. Thanks to this experiment, we could measure the weight of water remaining on the prosthesis as a function of the number of licks performed by the mouse following entry of the prosthesis on the water tank, and before retraction of the prosthesis (Supplementary Figure 1L). This analysis showed that substantial water was available on the prosthesis after exit from the water tank (approximately 10 mg), and that it was collected by the mouse in approximately 20 licks, after which there was almost no water left on the prosthesis. This confirms the relevance of the 20-lick threshold that we selected earlier.

### Simulation of the prosthesis

To perform controls while limiting animal experiments and minimizing prosthesis wear, we conceived a neural network model based on Long Short Term Memory (LSTM) layers (Supplementary Figure 2A). The network simulated the position of the prosthesis from spike trains as inputs (Supplementary Figure 2B), thus allowing to test behavioral performance that would result from spike trains that differ from the ones that actually took place. At each time step, the neuronal network calculated the trajectory of the prosthesis from the ongoing spike patterns it received, using the session-specific calibration values to translate spikes into prosthesis commands, that were then translated into simulated prosthesis movements by the neural network. To this aim, the neural network used the last 200 commands (to take into account prosthesis hysteresis), and an aging indicator of the prosthesis (Supplementary Figure 2A). The aging indicator was the normalized training day; it was necessary in order to take into account the wear of the prosthesis.

We validated this simulation method by measuring the deviation of simulated trajectories compared to real trajectories, by simulating all first and last sessions used for the analyses of Figures 3 and 4 (Supplementary Figure 2D-H). The deviation was sufficiently low for each dimension (Dim 1: 1.36%, Dim 2: 1.57%, Supplementary Figure 2C). When performing our control analysis, we altered the spike trains of the neurons and computed behavioral performance measures, such as the number of trajectories along the Main path (Figure 3M,O,P). Alterations were either a temporal shift between Dim 1 and Dim 2 spike trains (Supplementary Figure 2I, example with a 5 seconds shift on Dim 2 for visualization) or shuffling using a variable size of block. The shuffle was performed independently on Dim 1 and Dim 2 (Supplementary Figure 2J,K, examples with a 50 and 500 ms blocks). In Figures 3 and 4, shuffle data correspond to the average of 10 simulations. Note that for one session, we took advantage of this model to replace the tracking of the prosthesis that was lost for technical reasons. We were not able to apply the model on the 3D control space because of the lack of data to train the model accurately.

### Representation of averaged trajectory

To represent the averaged shape of multiple 2D trajectories (Figure 1D,E,G,I,J,L,N and Figure 5D), we used the dynamic time wrapping barycenter as defined in (Cuturi & Blondel, 2017) and implemented in the Tslearn python library (https://tslearn.readthedocs.io/en/stable/), using default parameters and gamma = 0.5.

### Pattern and trajectory detections

In this section, 1D patterns of firing rates and 1D trajectories are considered as the same mathematical object: a time series (TS) that repeats several times along the timeline of a behavioral session. In particular, for the analysis of trajectory and firing rate coordination (Figure 4), we needed to be able to identify the average shape of TS from a set of TSs (input TSs) aligned on the obtained rewards, and then find all similar TSs in shape across an entire session (output TSs), independently of the rewards. Towards this aim, we followed the following pipeline: (1) temporally align the input TSs that are the input of this algorithm; (2) characterize the shape of the input TSs and its variability through feature thresholds and finally (3) find all TSs in the session that verify those features that are returned as the output of this algorithm (output TSs).

(1) To accurately temporally align the input TSs, we first measured the average time course of those TSs without alignment. Then we searched the shift that maximized the Pearson correlation of each TS to this non-aligned mean. The mean of all aligned TSs became the average reference TS. Note that the input TSs could have different time duration. Thus, the average reference TS duration was set to the median of TS durations.
(2) We computed 4 features from the aligned TSs: Amplitude, defined as the difference between the max and the min of the TS, Euclidean distance to the reference, Pearson correlation to the reference, and Total variation of the TS defined as the sum of absolute differences between consecutive values.
(3) To detect all TS that matched the average reference TS in shape, we used a sliding window to scan the full time-course of the training sessions, and we selected the TS for which the 4 computer features fell within the 95% of the aligned TSs variability: meaning the 5^th^ percentile of the amplitudes and correlation distributions, the 95^th^ percentile of Euclidean distances distribution, and between the 2.5^th^ and 97.5^th^ of the total variation. This new set of TS was returned by the algorithm as the set of TSs that matched in shape that input TSs.

To sum up, this algorithm allowed us, from the set of 1D trajectory (or related 1D pattern of activity) corresponding to the Main path before a Lick Burst, to find all similar trajectories (or patterns of activity) independently of the Licks within a session.

### Measurement of coordination

To measure the coordination between the two set of 1D TSs (either 1D patterns or 1D trajectories) extracted separately from the two dimensions of the control space, we used two quantifications: (1) a coefficient of overlap, the Jaccard index, considering the time coverture of TS and (2) a coefficient of synchrony, the Spike Time Tiling Coefficient (STTC) computed on the TS onsets.

(1) The Jaccard index was defined as:

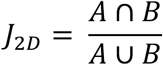

with A and B the time span of the TS from respectively Dimensions 1 and Dimensions 2. It can be interpreted as the amount of time when dimension 1 TS and dimension 2 TS take place simultaneously, normalized by the total time when one or the other TS take place. 0 means no overlapping while 1 means a perfect overlapping between A and B. In 3D, with A, B and C, the time span of the TS from respectively Dimensions 1, 2 and 3, the Jaccard index was defined equivalently as:

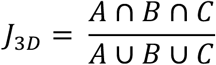
(2) STTC was applied in 2D as defined in Cutts & Eglen, 2014, considering the onset of 1D TS as an event (A and B for Dimension 1 and 2):

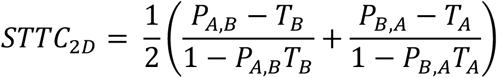

With *P_A,B_* the proportion of events A that occurs in a ±dt window around an event B, and vice-versa for *P_A,B_*, *T_A_* and *T_B_* are the total time covered by all the windows around respectively events A and B. STTC has been specifically designed to be robust to large differences of frequency between the two series of events. This coefficient measures the synchrony between A and B: 1 means that A events occurred within a chosen time window around B and vice-versa (synchronous activation). Conversely, –1 means that B occurs systematically outside of window around B and vice-versa (asynchronous activation). For our analyses, the selected window dt was set to ±500 ms around TS onsets. To apply the STTC in 3D, we extended the formula to take in account synchrony between each event to both others:

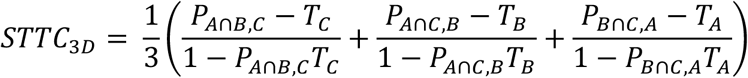

Where *P_A_*_∩*B,C*_ is the proportion of *A* ∩ *B* events that occurs in a ±dt window around an event C. *A* ∩ *B* events are all events of A that occur in a ±dt window around an event B combined with all events of B that occur in a ±dt window around an event A, i.e. all the synchronous events between A and B. An STTC of 1 means that all events A (and equivalently for B and C) occur within a ±dt window of an event B or C that is also within a ±dt window of an event C or B respectively.

## Acknowledgments

Experimental assistance and technical expertise were provided by Guillaume Hucher. We thank Marie Engel for work on an earlier version of the prosthesis, and Isabelle Ferezou for her constant support and comments on the manuscript.

This work was funded by CNRS (CNRS 80|Prime, PRIME interdisciplinary label), Fondation pour la Recherche Médicale, La Fondation Dassault Systèmes, Agence Nationale pour la Recherche (ANR JCJC Mesobrain, ANR PRC Expect, Motorsense, Hermin), Lidex NeuroSaclay, Idex Brainscopes, iCODE, hCODE.

---

**Supplementary Figure 1:**
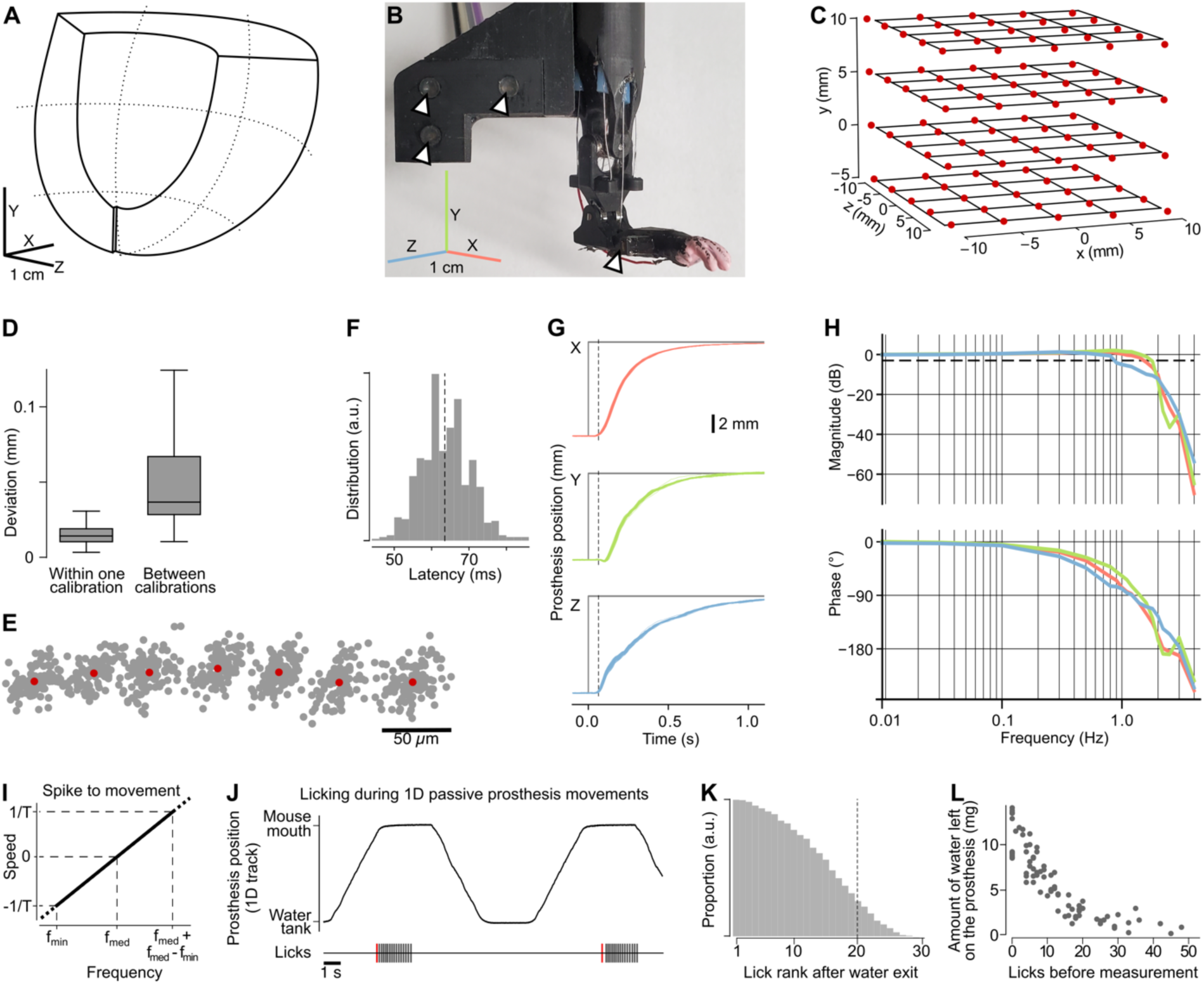
Mechanical and task-related characterization of the prosthesis. (**A**) Volume that can be reached by the end effector of the prosthesis. Within this volume, the prosthesis can be controlled in cartesian coordinates (X,Y,Z coordinates shown next to the volume), or in coordinates that are specific to the ongoing task, such as the 2D subspace of Figure 1D. (**B**) Prosthesis skeleton featuring the LED landmarks for calibration of the tracking system, as well as the LED located on the prosthesis end effector (arrows). (**C**) Displacement of the prosthesis LED tracked by the cameras (red dots) compared to displacements imposed with a 3D linear translation stage rated for micrometric precision (black grid). Each red dot is the mean of 100 measures. (**D**) Variation of the LED position within one landmark calibration (100 measures, mean = 15 um) and between 20 landmark calibrations (mean of 100 measures for each calibration, mean = 50 um). (**E**) Position of the prosthesis LED with controlled displacements (using one axis of the 3D micrometric linear stage) of 50 µm steps. Grey dots are individual measures and black dots are the mean position measured at each step. (**F**) Histogram of latencies between 10 mm step commands and the observed movement onsets. Dashed line: mean latency 63.5 ms. (**G**) Superimposed 100 responses to a 10 mm step command, in each of the 3 dimensions of the cartesian control space (X,Y and Z). Dashed lines are the mean latencies. (**H**) Bode plots measured on prosthesis movements in response to sinusoidal commands of varying frequencies. Red, green and blue: movements following the X, Y and Z command axes. Top: magnitude of the prosthesis movements. Bottom: dephasing of the prosthesis movements. Dashed line? (**I**) Conversion of the firing rate of a neuron into a speed command. Firing rates were linearly converted to speed using a formula based on f_min_, the 1^st^ percentile of the baseline activity, and f_med_ the median of the baseline activity. See methods. (**J**) Top: automated movements of the prosthesis in Dimension 1 from the water tank to the mouse mouth area during habituation. Bottom: bursts of licks performed by the mouse on the prosthesis. Red: first lick after an exit of the prosthesis from the water tank and start of the water reward consumption. (**K**) Distribution of the position of licks with respect to the last Water trip, in the bursts measured across 2 mice (4 sessions in total) during passive, automated water to mouth trajectories. Dashed line: 95^th^ percentile, corresponding to 20 licks. This average lick count is used in the analysis as the estimated maximum number of licks that result in water intake during active control of the prosthesis. (**L**) Weight of the water left on the prosthesis after it is plunged on the water tank and was licked by a mouse (Y axis), as function of the total number of licks performed by the mouse from water tank exit until water weight measurement (X axis).

**Supplementary Figure 2:**
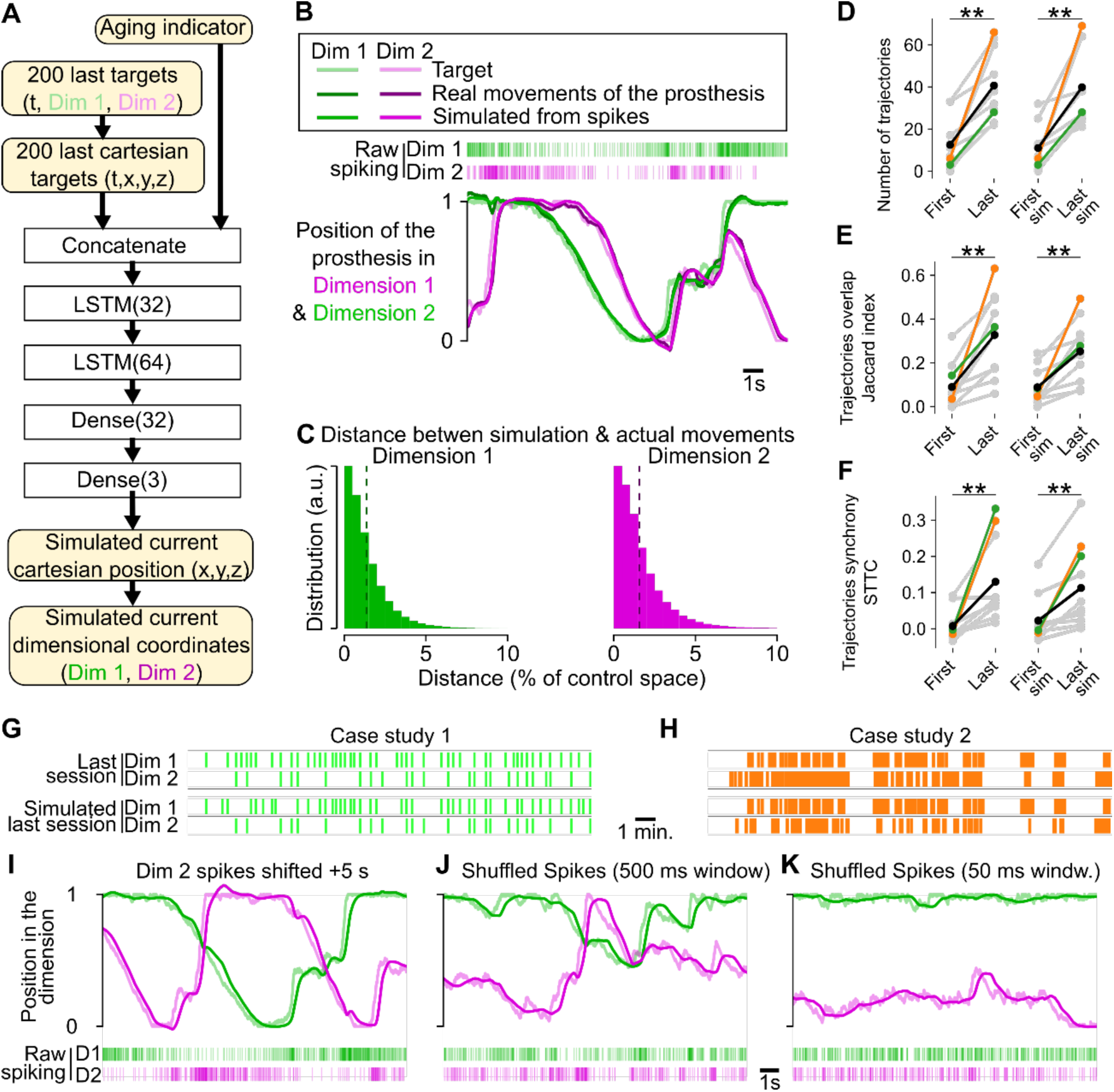
Simulation of the prosthesis movements. (**A**) Diagram of the recurrent neuronal network used to simulate the movements of the prosthesis for simulations. **(B)** Example of simulated prosthesis movements, compared to the actual prosthesis movements triggered by the corresponding target. **(C)** Distribution of the distance between the position predicted by the model from the target, and the actual positions of the prosthesis in response to the target. The distance is a % of the control space. Dashed line denotes means: 1.36% (Dim 1) and 1.57% (Dim 2). **(D)** Number of 2D Main trajectories (see Figure 3) between the first and last training session in Learning sequences. Left graph: observed prosthesis movements. Right graph: simulated movements. Orange and green lines: Case Studies 1 and 2. **: Wilcoxon test, p = 0.002. **(E)** Same as **D** for the Jaccard index of overlap between the 1D Main patterns measured in dimensions 1 and 2. **: Wilcoxon test, p = 0.002. **(F)** Same as **E** for an STTC estimate of synchrony. **: Wilcoxon test, p = 0.006. **(G)** Prosthesis 1D Main trajectories in Case study 1. The timing of Dimensions 1 and 2 of the Main trajectories (see Figure 4) that were recorded in the actual prosthesis movements (top, green) match the timing in the simulated prosthesis trajectories (bottom, black). **(H)** Same as **G** for Case Study 2. **(I)** Example of the impact of a time shift between the two control neurons on the target and on the simulated prosthesis movements. **(J)** Same as I for a shuffle with a 500 ms permutation block. **(K)** Same as I for a shuffle with a 50 ms permutation block.

## Bibliography

1. Aach, M., Schildhauer, T. A., Zieriacks, A., Jansen, O., Weßling, M., Brinkemper, A., & Grasmücke, D. (2023). Feasibility, safety, and functional outcomes using the neurological controlled Hybrid Assistive Limb exoskeleton (HAL®) following acute incomplete and complete spinal cord injury – Results of 50 patients. The Journal of Spinal Cord Medicine, 46(4), 574–581. 10.1080/10790268.2023.2200362

2. Abbasi, A., Goueytes, D., Shulz, D. E., Ego-Stengel, V., & Estebanez, L. (2018). A fast intracortical brain– machine interface with patterned optogenetic feedback. Journal of Neural Engineering, 15(4), 046011. 10.1088/1741-2552/aabb80

3. Abbasi, A., Lassagne, H., Estebanez, L., Goueytes, D., Shulz, D. E., & Ego-Stengel, V. (2023). Brain-machine interface learning is facilitated by specific patterning of distributed cortical feedback. SCIENCE ADVANCES.

4. Alonso, I., Scheer, I., Palacio-Manzano, M., Frézel-Jacob, N., Philippides, A., & Prsa, M. (2023). Peripersonal encoding of forelimb proprioception in the mouse somatosensory cortex. Nature Communications, 14(1), 1866. 10.1038/s41467-023-37575-w

5. Arduin, P.-J., Frégnac, Y., Shulz, D. E., & Ego-Stengel, V. (2013). “Master” Neurons Induced by Operant Conditioning in Rat Motor Cortex during a Brain-Machine Interface Task. The Journal of Neuroscience, 33(19), 8308–8320. 10.1523/JNEUROSCI.2744-12.2013

6. Arduin, P.-J., Fregnac, Y., Shulz, D. E., & Ego-Stengel, V. (2014). Bidirectional control of a one-dimensional robotic actuator by operant conditioning of a single unit in rat motor cortex. Frontiers in Neuroscience, 8. 10.3389/fnins.2014.00206

7. Bach Baunsgaard, C., Vig Nissen, U., Katrin Brust, A., Frotzler, A., Ribeill, C., Kalke, Y.-B., León, N., Gómez, B., Samuelsson, K., Antepohl, W., Holmström, U., Marklund, N., Glott, T., Opheim, A., Benito, J., Murillo, N., Nachtegaal, J., Faber, W., & Biering-Sørensen, F. (2018). Gait training after spinal cord injury: Safety, feasibility and gait function following 8 weeks of training with the exoskeletons from Ekso Bionics. Spinal Cord, 56(2), 106–116. 10.1038/s41393-017-0013-7

8. Barrett, J. M., Martin, M. E., & Shepherd, G. M. G. (2022). Manipulation-specific cortical activity as mice handle food. Current Biology, 32(22), 4842–4853.e6. 10.1016/j.cub.2022.09.045

9. Barrett, J. M., Raineri Tapies, M. G., & Shepherd, G. M. G. (2020). Manual dexterity of mice during food-handling involves the thumb and a set of fast basic movements. PLOS ONE, 15(1), e0226774. 10.1371/journal.pone.0226774

10. Carmena, J. M., Lebedev, M. A., Crist, R. E., O’Doherty, J. E., Santucci, D. M., Dimitrov, D. F., Patil, P. G., Henriquez, C. S., & Nicolelis, M. A. L. (2003). Learning to Control a Brain–Machine Interface for Reaching and Grasping by Primates. PLoS Biology, 1(2), e42. 10.1371/journal.pbio.0000042

11. Carmena, J. M., Lebedev, M. A., Henriquez, C. S., & Nicolelis, M. A. L. (2005). Stable ensemble performance with single-neuron variability during reaching movements in primates. Journal of Neuroscience, 25(46), 10712–10716. 10.1523/JNEUROSCI.2772-05.2005

12. Chapin, J. K., Moxon, K. A., Markowitz, R. S., & Nicolelis, M. A. L. (1999). Real-time control of a robot arm using simultaneously recorded neurons in the motor cortex. Nature Neuroscience, 2(7), 664–670. 10.1038/10223

13. Clancy, K. B., Koralek, A. C., Costa, R. M., Feldman, D. E., & Carmena, J. M. (2014). Volitional modulation of optically recorded calcium signals during neuroprosthetic learning. Nature Neuroscience, 17(6), 807–809. 10.1038/nn.3712

14. Clancy, K. B., & Mrsic-Flogel, T. D. (2021). The sensory representation of causally controlled objects. Neuron, 109(4), 677–689.e4. 10.1016/j.neuron.2020.12.001

15. Cutts, C. S., & Eglen, S. J. (2014). Detecting Pairwise Correlations in Spike Trains: An Objective Comparison of Methods and Application to the Study of Retinal Waves. The Journal of Neuroscience, 34(43), 14288–14303. 10.1523/JNEUROSCI.2767-14.2014

16. Cuturi, M., & Blondel, M. (2017). Soft-DTW: a Differentiable Loss Function for Time-Series. Proceedings of the 34 Th International Conference on Machine Learning.

17. del-Ama, A. J., Gil-Agudo, Á., Pons, J. L., & Moreno, J. C. (2014). Hybrid FES-robot cooperative control of ambulatory gait rehabilitation exoskeleton. Journal of NeuroEngineering and Rehabilitation, 11(1), 27. 10.1186/1743-0003-11-27

18. Dogadov, A. A., Shulz, D. E., Ferezou, I., Ego-Stengel, V., & Estebanez, L. (2024). Operant conditioning of cortical waves through a brain-machine interface. 10.1101/2024.09.10.612344

19. Estebanez, L., Hoffmann, D., Voigt, B. C., & Poulet, J. F. A. (2017). Parvalbumin-Expressing GABAergic Neurons in Primary Motor Cortex Signal Reaching. Cell Reports, 20(2), 308–318. 10.1016/j.celrep.2017.06.044

20. Fajardo, J., Ferman, V., Cardona, D., Maldonado, G., Lemus, A., & Rohmer, E. (2020). Galileo Hand: An Anthropomorphic and Affordable Upper-Limb Prosthesis. IEEE Access, 8, 81365–81377. 10.1109/ACCESS.2020.2990881

21. Fetz, E. E. (1969). Operant Conditioning of Cortical Unit Activity.

22. Flesher, S. N., Downey, J. E., Weiss, J. M., Hughes, C. L., Herrera, A. J., Tyler-Kabara, E. C., Boninger, M. L., Collinger, J. L., & Gaunt, R. A. (2021). A brain-computer interface that evokes tactile sensations improves robotic arm control. Science, 372(6544), 831–836. 10.1126/science.abd0380

23. Fox, J. G. (2007). The mouse in biomedical research (2nd ed). Academic press.

24. Galiñanes, G. L., Bonardi, C., & Huber, D. (2018). Directional Reaching for Water as a Cortex-Dependent Behavioral Framework for Mice. Cell Reports, 22(10), 2767–2783. 10.1016/j.celrep.2018.02.042

25. Georgopoulos, A., Kalaska, J., Caminiti, R., & Massey, J. (1982). On the relations between the direction of two-dimensional arm movements and cell discharge in primate motor cortex. The Journal of Neuroscience, 2(11), 1527–1537. 10.1523/JNEUROSCI.02-11-01527.1982

26. Golub, M. D., Sadtler, P. T., Oby, E. R., Quick, K. M., Ryu, S. I., Tyler-Kabara, E. C., Batista, A. P., Chase, S. M., & Yu, B. M. (2018). Learning by neural reassociation. Nature Neuroscience, 21(4), 607–616. 10.1038/s41593-018-0095-3

27. Goueytes, D., Abbasi, A., Lassagne, H., Shulz, D. E., Estebanez, L., & Ego-Stengel, V. (2019). Control of a robotic prosthesis simulation by a closed-loop intracortical brain-machine interface. 2019 9th International IEEE/EMBS Conference on Neural Engineering (NER), 183–186. 10.1109/NER.2019.8716885

28. Goueytes, D., Lassagne, H., Shulz, D. E., Ego-Stengel, V., & Estebanez, L. (2022). Learning in a closed-loop brain-machine interface with distributed optogenetic cortical feedback. Journal of Neural Engineering, 19(6), 066045. 10.1088/1741-2552/acab87

29. Guo, J.-Z., Graves, A. R., Guo, W. W., Zheng, J., Lee, A., Rodríguez-González, J., Li, N., Macklin, J. J., Phillips, J. W., Mensh, B. D., Branson, K., & Hantman, A. W. (2015). Cortex commands the performance of skilled movement. eLife, 4, e10774. 10.7554/eLife.10774

30. Guo, K., Yamawaki, N., Barrett, J. M., Tapies, M., & Shepherd, G. M. G. (2020). Cortico-Thalamo-Cortical Circuits of Mouse Forelimb S1 Are Organized Primarily as Recurrent Loops. The Journal of Neuroscience, 40(14), 2849– 2858. 10.1523/JNEUROSCI.2277-19.2020

31. Hayatou, Z., Wang, H., Chaillet, A., Shulz, D. E., Ego-Stengel, V., & Estebanez, L. (2025). Embodiment of an artificial limb in mice. PLOS Biology, 23(6), e3003186. 10.1371/journal.pbio.3003186

32. Hochberg, L. R., Bacher, D., Jarosiewicz, B., Masse, N. Y., Simeral, J. D., Vogel, J., Haddadin, S., Liu, J., Cash, S. S., van der Smagt, P., & Donoghue, J. P. (2012). Reach and grasp by people with tetraplegia using a neurally controlled robotic arm. Nature, 485(7398), 372–375. 10.1038/nature11076

33. Ifft, P. J., Shokur, S., Li, Z., Lebedev, M. A., & Nicolelis, M. A. L. (2013). A Brain-Machine Interface Enables Bimanual Arm Movements in Monkeys. Science Translational Medicine, 5(210). 10.1126/scitranslmed.3006159

34. Ijspeert, A. J., Crespi, A., Ryczko, D., & Cabelguen, J.-M. (2007). From Swimming to Walking with a Salamander Robot Driven by a Spinal Cord Model. Science, 315(5817), 1416–1420. 10.1126/science.1138353

35. Johannes, M. S., Faulring, E. L., Katyal, K. D., Para, M. P., Helder, J. B., Makhlin, A., Moyer, T., Wahl, D., Solberg, J., Clark, S., Armiger, R. S., Lontz, T., Geberth, K., Moran, C. W., Wester, B. A., Van Doren, T., & Santos-Munne, J. J. (2020). The Modular Prosthetic Limb. In Wearable Robotics (pp. 393–444). Elsevier. 10.1016/B978-0-12-814659-0.00021-7

36. Karpowicz, B. M., Ali, Y. H., Wimalasena, L. N., Sedler, A. R., Keshtkaran, M. R., Bodkin, K., Ma, X., Rubin, D. B., Williams, Z. M., Cash, S. S., Hochberg, L. R., Miller, L. E., & Pandarinath, C. (2025). Stabilizing brain-computer interfaces through alignment of latent dynamics. Nature Communications, 16(1), 4662. 10.1038/s41467-025-59652-y

37. Kerdraon, J., Previnaire, J. G., Tucker, M., Coignard, P., Allegre, W., Knappen, E., & Ames, A. (2021). Evaluation of safety and performance of the self balancing walking system Atalante in patients with complete motor spinal cord injury. Spinal Cord Series and Cases, 7(1), 71. 10.1038/s41394-021-00432-3

38. Keysers, C., & Michon, F. (2024). Can mirror self-recognition in mice unpack the neural underpinnings of self-awareness? Neuron, 112(2), 177–179. 10.1016/j.neuron.2023.12.005

39. Koralek, A. C., Jin, X., Long Ii, J. D., Costa, R. M., & Carmena, J. M. (2012). Corticostriatal plasticity is necessary for learning intentional neuroprosthetic skills. Nature, 483(7389), 331–335. 10.1038/nature10845

40. Le, H. M., Do, T. N., & Phee, S. J. (2016). A survey on actuators-driven surgical robots. Sensors and Actuators A: Physical, 247, 323–354. 10.1016/j.sna.2016.06.010

41. Moritz, C. T., Perlmutter, S. I., & Fetz, E. E. (2008). Direct control of paralysed muscles by cortical neurons. Nature, 456(7222), 639–642. 10.1038/nature07418

42. Oby, E. R., Degenhart, A. D., Grigsby, E. M., Motiwala, A., McClain, N. T., Marino, P. J., Yu, B. M., & Batista, A. P. (2025). Dynamical constraints on neural population activity. Nature Neuroscience, 28(2), 383–393. 10.1038/s41593-024-01845-7

43. Oby, E. R., Golub, M. D., Hennig, J. A., Degenhart, A. D., Tyler-Kabara, E. C., Yu, B. M., Chase, S. M., & Batista, A. P. (2019). New neural activity patterns emerge with long-term learning. Proceedings of the National Academy of Sciences of the United States of America, 116(30), 15210–15215. 10.1073/pnas.1820296116

44. Orsborn, A. L., Moorman, H. G., Overduin, S. A., Shanechi, M. M., Dimitrov, D. F., & Carmena, J. M. (2014). Closed-loop decoder adaptation shapes neural plasticity for skillful neuroprosthetic control. Neuron, 82(6), 1380– 1393. 10.1016/j.neuron.2014.04.048

45. Pasquini, M., James, N. D., Dewany, I., Coen, F.-V., Cho, N., Lai, S., Anil, S., Carpaneto, J., Barraud, Q., Lacour, S. P., Micera, S., & Courtine, G. (2022). Preclinical upper limb neurorobotic platform to assess, rehabilitate, and develop therapies. Science Robotics, 7(64), eabk2378. 10.1126/scirobotics.abk2378

46. Prescott, T. J., Vogeley, K., & Wykowska, A. (2024). Understanding the sense of self through robotics. Science Robotics, 9(95), eadn2733. 10.1126/scirobotics.adn2733

47. Prsa, M., Galiñanes, G. L., & Huber, D. (2017). Rapid Integration of Artificial Sensory Feedback during Operant Conditioning of Motor Cortex Neurons. Neuron, 93(4), 929–939.e6. 10.1016/j.neuron.2017.01.023

48. Pyasik, M., Tieri, G., & Pia, L. (2020). Visual appearance of the virtual hand affects embodiment in the virtual hand illusion. Scientific Reports, 10(1), 5412. 10.1038/s41598-020-62394-0

49. Raspopovic, S., Capogrosso, M., Petrini, F. M., Bonizzato, M., Rigosa, J., Di Pino, G., Carpaneto, J., Controzzi, M., Boretius, T., Fernandez, E., Granata, G., Oddo, C. M., Citi, L., Ciancio, A. L., Cipriani, C., Carrozza, M. C., Jensen, W., Guglielmelli, E., Stieglitz, T., … Micera, S. (2014). Restoring Natural Sensory Feedback in Real-Time Bidirectional Hand Prostheses. Science Translational Medicine, 6(222). 10.1126/scitranslmed.3006820

50. Resnik, L., Klinger, S. L., & Etter, K. (2014). The DEKA Arm: Its features, functionality, and evolution during the Veterans Affairs Study to optimize the DEKA Arm. Prosthetics & Orthotics International, 38(6), 492–504. 10.1177/0309364613506913

51. Risso, G., Preatoni, G., Valle, G., Marazzi, M., Bracher, N. M., & Raspopovic, S. (2022). Multisensory stimulation decreases phantom limb distortions and is optimally integrated. iScience, 25(4), 104129. 10.1016/j.isci.2022.104129

52. Rognini, G., Petrini, F. M., Raspopovic, S., Valle, G., Granata, G., Strauss, I., Solcà, M., Bello-Ruiz, J., Herbelin, B., Mange, R., D’Anna, E., Di Iorio, R., Di Pino, G., Andreu, D., Guiraud, D., Stieglitz, T., Rossini, P. M., Serino, A., Micera, S., & Blanke, O. (2019). Multisensory bionic limb to achieve prosthesis embodiment and reduce distorted phantom limb perceptions. *Journal of Neurology*, Neurosurgery & Psychiatry, 90(7), 833–836. 10.1136/jnnp-2018-318570

53. Sadtler, P. T., Quick, K. M., Golub, M. D., Chase, S. M., Ryu, S. I., Tyler-Kabara, E. C., Yu, B. M., & Batista, A. P. (2014). Neural constraints on learning. Nature, 512(7515), 423–426. 10.1038/nature13665

54. Sorrell, E., Rule, M. E., & O’Leary, T. (2021). Brain–Machine Interfaces: Closed-Loop Control in an Adaptive System. *Annual Review of Control*, Robotics, and Autonomous Systems, 4(1), 167–189. 10.1146/annurev-control-061720-012348

55. Tieri, G., Gioia, A., Scandola, M., Pavone, E. F., & Aglioti, S. M. (2017). Visual appearance of a virtual upper limb modulates the temperature of the real hand: A thermal imaging study in Immersive Virtual Reality. European Journal of Neuroscience, 45(9), 1141–1151. 10.1111/ejn.13545

56. Umezawa, K., Suzuki, Y., Ganesh, G., & Miyawaki, Y. (2022). Bodily ownership of an independent supernumerary limb: An exploratory study. Scientific Reports, 12(1), 2339. 10.1038/s41598-022-06040-x

57. Valle, G., Alamri, A. H., Downey, J. E., Lienkämper, R., Jordan, P. M., Sobinov, A. R., Endsley, L. J., Prasad, D., Boninger, M. L., Collinger, J. L., Warnke, P. C., Hatsopoulos, N. G., Miller, L. E., Gaunt, R. A., Greenspon, C. M., & Bensmaia, S. J. (2025). Tactile edges and motion via patterned microstimulation of the human somatosensory cortex. Science, 387(6731), 315–322. 10.1126/science.adq5978

58. Van Den Brand, R., Heutschi, J., Barraud, Q., DiGiovanna, J., Bartholdi, K., Huerlimann, M., Friedli, L., Vollenweider, I., Moraud, E. M., Duis, S., Dominici, N., Micera, S., Musienko, P., & Courtine, G. (2012). Restoring Voluntary Control of Locomotion after Paralyzing Spinal Cord Injury. Science, 336(6085), 1182–1185. 10.1126/science.1217416

59. Velliste, M., Perel, S., Spalding, M. C., Whitford, A. S., & Schwartz, A. B. (2008). Cortical control of a prosthetic arm for self-feeding. Nature, 453(7198), 1098–1101. 10.1038/nature06996

60. Vestergaard, M., Carta, M., Güney, G., & Poulet, J. F. A. (2023). The cellular coding of temperature in the mammalian cortex. Nature, 614(7949), 725–731. 10.1038/s41586-023-05705-5

61. Vigaru, B. C., Lambercy, O., Schubring-Giese, M., Hosp, J. A., Schneider, M., Osei-Atiemo, C., Luft, A., & Gassert, R. (2013). A Robotic Platform to Assess, Guide and Perturb Rat Forelimb Movements. IEEE Transactions on Neural Systems and Rehabilitation Engineering, 21(5), 796–805. 10.1109/TNSRE.2013.2240014

62. Wada, M., Takano, K., Ora, H., Ide, M., & Kansaku, K. (2016). The Rubber Tail Illusion as Evidence of Body Ownership in Mice. The Journal of Neuroscience, 36(43), 11133–11137. 10.1523/JNEUROSCI.3006-15.2016

63. Wagner, F. B., Mignardot, J.-B., Le Goff-Mignardot, C. G., Demesmaeker, R., Komi, S., Capogrosso, M., Rowald, A., Seáñez, I., Caban, M., Pirondini, E., Vat, M., McCracken, L. A., Heimgartner, R., Fodor, I., Watrin, A., Seguin, P., Paoles, E., Van Den Keybus, K., Eberle, G., … Courtine, G. (2018). Targeted neurotechnology restores walking in humans with spinal cord injury. Nature, 563(7729), 65–71. 10.1038/s41586-018-0649-2

64. Wang, H., Stout, D. B., & Chatziioannou, A. F. (2015). A Deformable Atlas of the Laboratory Mouse. Molecular Imaging and Biology, 17(1), 18–28. 10.1007/s11307-014-0767-7

65. Wodlinger, B., Downey, J. E., Tyler-Kabara, E. C., Schwartz, A. B., Boninger, M. L., & Collinger, J. L. (2015). Ten-dimensional anthropomorphic arm control in a human brain−machine interface: Difficulties, solutions, and limitations. Journal of Neural Engineering, 12(1), 016011. 10.1088/1741-2560/12/1/016011

66. Yamawaki, N., Raineri Tapies, M. G., Stults, A., Smith, G. A., & Shepherd, G. M. (2021). Circuit organization of the excitatory sensorimotor loop through hand/forelimb S1 and M1. eLife, 10, e66836. 10.7554/eLife.66836

67. Yang, W., Kanodia, H., & Arber, S. (2023). Structural and functional map for forelimb movement phases between cortex and medulla. Cell, 186(1), 162–177.e18. 10.1016/j.cell.2022.12.009

